# Dia2 formin controls receptor activity by organizing plasma membrane lipid partitioning at the nanoscale

**DOI:** 10.1101/2023.12.15.571857

**Authors:** Changting Li, Yannick Hamon, David Mazaud, Pamela Gonzalez Troncoso, Marie Dessard, Hai-tao He, Christophe Lamaze, Cedric M. Blouin

## Abstract

Activation of JAK/STAT signaling by IFN-γ requires partitioning of IFN-γR into specific lipid nanodomains at the plasma membrane. Using IFN-γR as a proxy, we investigated the role of actin dynamics in the formation and organization of lipid nanodomains, a process that remains poorly understood. We identified formin Dia2/DIAPH3 as a specific and RhoA -dependent regulator of IFN-γ-induced JAK/STAT signaling. Based on lipidomics and specific probes enabling membrane lipid imaging by super resolution microscopy, we demonstrate that Dia2 is required for proper assembly of sphingomyelin and cholesterol lipid complexes. Finally, we show that the disorganization of lipid nanodomains induced by Dia2 depletion results in drastic changes in nano-partitioning and activity of other membrane proteins, such as Thy1 and PD-L1. Our data establish, therefore, the central role of the RhoA-Dia2 axis in the regulation of IFN-γ induced JAK/STAT signaling, and more broadly, in the nanoscale organization of the plasma membrane.

## Introduction

The spatio-temporal distribution and dynamics of lipids at the plasma membrane condition the lateral segregation of signaing receptors into functional nanodomains (Rao & Mayor, 2014; Sezgin et al., 2017) that play a key role in their activation (Levental & Lyman, 2023). Receptor nano-partitioning is required for activation of T cell and B cell receptors (Gold & Reth, 2019; Pathan-Chhatbar et al., 2021), interferon-γ receptor (IFN-γR) (Blouin et al., 2016), GPCRs (Baccouch et al., 2022), integrins (Kalappurakkal et al., 2019), and controls various cellular processes such as immune response, cell adhesion and cell survival (Saha et al., 2016). Since the concept of lipid rafts was first proposed (Simons & Ikonen, 1997), there have been persistent controversies about the nature and function of the physical constraints that control the spatiotemporal diffusion and biological activity of plasma membrane components (Sezgin et al., 2017; Levental et al., 2020). Importantly, new technological developments, such as super-resolution microscopy, enhanced mass spectrometry and molecular dynamics simulations have led to a recent revision of the original concept. Lipid rafts are now considered as highly dynamic nanoscale lipid-protein assemblies enriched with cholesterol and sphingolipids, and where receptors such as glycosylphosphatidylinositol-anchored proteins (GPI-APs) and cytokine transmembrane receptors are preferentially recruited (Levental et al., 2020; Murata et al., 2022). Besides that, the cortical actin cytoskeleton has been shown to influence membrane organization and determine molecular diffusion at the plasma membrane (Jaqaman et al. 2011; Köster et al., 2016; Fritzsche et al., 2017; Kusumi et al., 2023). The actin-driven diffusion mechanism of membrane molecules can provide a valid explanation for the kinetic properties and non-equilibrium distribution of nanoclusters formed by multiple lipid or protein species. Nevertheless, the current paradigm of ordered lipid nanodomains still leaves us with several processes that are poorly understood, such as the identification of drivers for interactions between lipids, receptors, and actin within membrane nanodomains, and the role of these interactions in receptor signal transduction and physiological functions.

We demonstrated previously that the IFN-γ receptor (IFN-γR) partitions into sphingomyelin (SM) and cholesterol (Chol)-dependent nanodomains, a process that is strictly required for IFN-γR conformational changes and subsequent Janus kinases/Signal Transducers and Activators of Transcription (JAK/STAT) signaling (Blouin et al., 2016). Given the significant role of actin cytoskeleton in membrane organization, we sought to understand how it may be involved in the association of IFN-γR with specific plasma membrane nanodomains and the regulation of JAK/STAT signaling. Formins have been shown to play a central role in the control of actin nucleation and elongation. They are involved in several cellular processes, including filopodia and lamellipodia formation (Schirenbeck et al., 2005, Block et al., 2012), cytokinesis (Buracco et al., 2019), and phagocytic cups formation (Brandt et al., 2007). The formin family groups 15 formin genes in placental mammals which can be categorized into seven subtypes (Courtemanche, 2018): Diaphanous (DIAPH/Dia), Dishevelled-Associated Activator of Morphogenesis (DAAM), Formin-Related gene in Leukocytes (FRL), Formin Homology Domain-containing Protein (FHOD), Inverted Formin (INF), Formin like (FMN) protein, and Delphilin (Kovar et al., 2005). The role of formins in actin nucleation and filament elongation has been extensively studied, but their specific role in the function of plasma membrane transmembrane receptors has not been extensively addressed. Recently, formins have been involved in the diffusion of GPI-APs and B cell receptor at the plasma membrane, (Saha et al., 2015; Kalappurakkal et al., 2019; Rey-Suarez et al., 2020) and in the formation of CD44 nanoclusters (Freeman et al., 2018; Sil et al., 2020). However, the precise molecular mechanisms by which formins drive the formation of specific lipid nanodomains and possibly control membrane receptor signaling remain largely elusive.

We identified DIAPH3/Dia2, a member of the formin family, as an essential and specific regulator of IFN-γ-stimulated JAK/STAT signaling. Spot-variation fluorescence correlation spectroscopy (svFCS) revealed a drastic change of membrane receptor diffusion behavior upon Dia2 inhibition. Spinning-disk and single molecule localization microscopy respectively showed that Dia2 depletion decreased the amount of cholesterol-associated sphingomyelin (SM/Chol complex) at the plasma membrane and its nanoclustering. Furthermore, we identified RhoA as a major upstream regulator of Dia2 controlling IFN-γR function. We next showed that the RhoA-Dia2 axis has a general impact on plasma membrane nano-organization, as evidenced by the perturbed diffusion of GPI-AP Thy-1 in absence of Dia2. Finally, we observed that Dia2 was important for both the cellular abundance and the protein stability of PD-L1, another receptor controlled by its lipid environment at the plasma membrane. Altogether, our data demonstrate that the RhoA-Dia2 axis governs the formation of SM/Chol nanodomains that are required for proper membrane receptors nano-partitioning and function at the plasma membrane.

## Results

### Formin inhibitor SMIFH2 impairs IFN-γR-induced JAK/STAT signaling

IFN-γ is a pleiotropic cytokine that regulates many cellular functions and plays a key role in innate immunity but also a dual function in tumor progression and elimination (Stark & Darnell, 2012; Castro et al., 2018). IFN-γR is a heterotetramer composed of two copies of IFN-γR1 and two copies of IFN-γR2 chains. Upon ligand binding, STATs are phosphorylated by receptor-docked JAKs and then dissociate from the receptor to rapidly translocate to the nucleus as transcription factors that bind IFN-γ-activated site (GAS) elements and induce IFN-stimulated genes (ISGs) activation (Stark & Darnell, 2012; Philips et al., 2022). To investigate the role of actin in IFN-γR signaling function, we first treated human immortalized fibroblasts with several specific actin-perturbing drugs prior to IFN-γ stimulation. Only the formin specific inhibitor, SMIFH2 decreased the tyrosine phosphorylation of STAT1 (pSTAT1) upon IFN-γ stimulation (Fig. 1a). We did not observe any difference for pSTAT1 levels in cells treated with other actin related inhibitors, such as the actin monomer sequestering drug Latrunculin A (LatA), the actin polymerization inhibitor Cytochalasin D (CytoD), Arp2/3 inhibitor CK666, and myosin-II inhibitor Blebbistatin (Fig. 1a). Moreover, we recently showed that SMIFH2 and its derived compounds can directly inhibit IFN-γ activity in the extracellular medium (Thoidingjam et al., 2022). To eliminate extracellular SMIFH2 before cytokine addition, we added a PBS washing step (SMI-WASH) prior to IFN-γ stimulation (Fig. 1b). We first verified that SMI-WASH condition still significantly reduces F-actin polymerization as visualized by phalloidin staining (Extended data Fig. 1a), consistent with its original inhibitory effect on formin-mediated actin elongation (Rizvi et al., 2009). We observed that SMI-WASH still significantly decreased STAT1 phosphorylation level (Fig. 1c) and nuclear translocation (Fig. 1d) without affecting IFN-γ binding and uptake (Extended data Fig. 1b). To further characterize the impact of formin inhibition on JAK/STAT signaling, we monitored the activation of JAK1 and JAK2 that are respectively associated with IFN-γR1 and IFN-γR2, and upstream of STAT1 phosphorylation. We found that SMI-WASH treatment decreased both JAK1 (Fig. 1e) and JAK2 (Fig. 1f) phosphorylation levels upon IFN-γ stimulation. All together, we concluded that formin inhibitor SMIFH2 specifically inhibits early stage of IFN-γ mediated JAK/STAT activation at the plasma membrane.

**Fig. 1.**
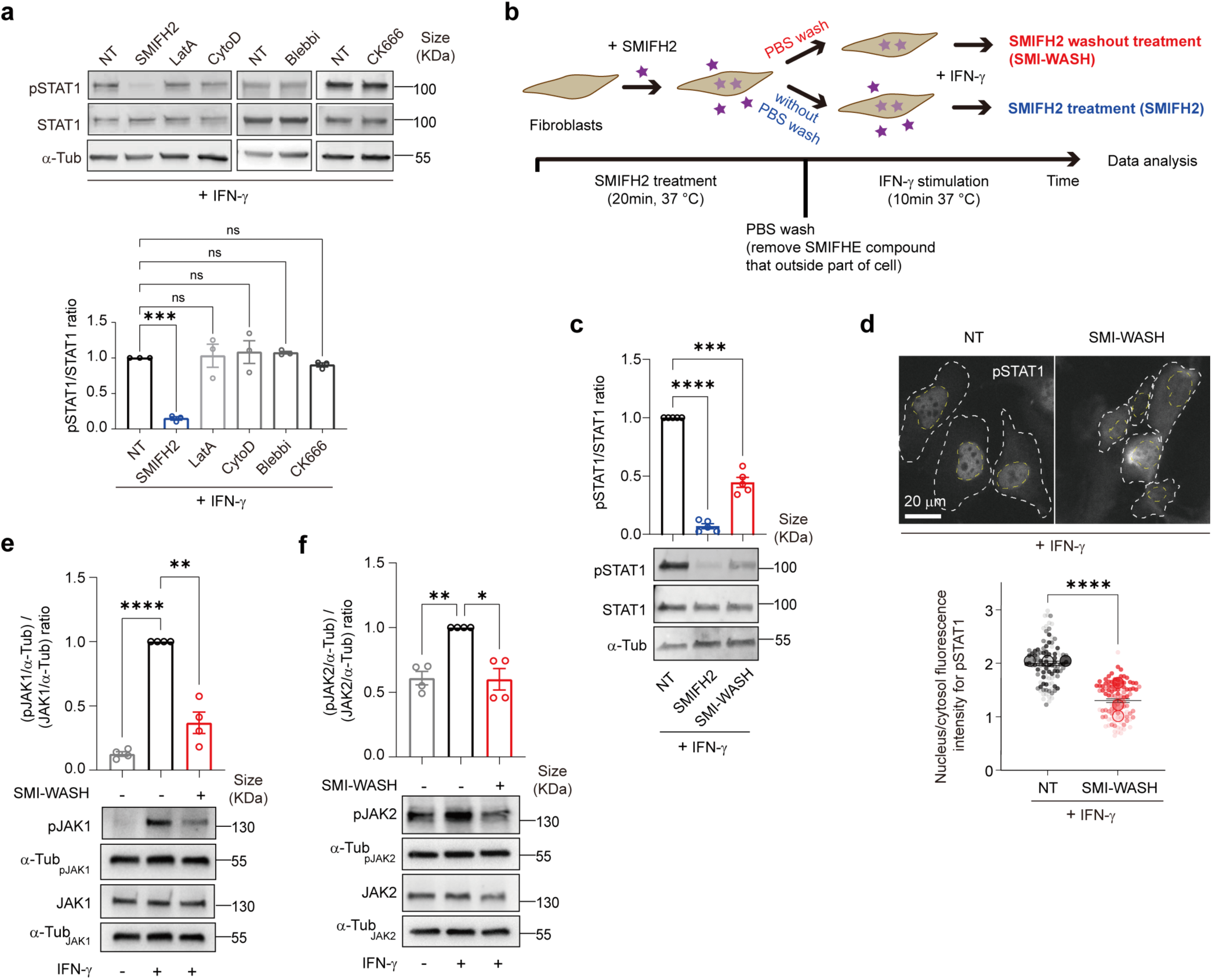
Formin inhibitor SMIFH2 impairs IFN-γR-induced JAK/STAT signaling. (**a**) Western blotting analysis for <u>tyrosine phosphorylation</u> status of STAT1 in cell lysates of fibroblasts stimulated for 10 min with IFN-γ (1000 U/ml) in presence of either SMIFH2 (25 μM), Latrunculin A (LatA) (500 nM), Cytochalasin D (Cyto D) (1 μM), CK666 (50 μM) or Blebbistation (25 μM). Quantification for phosphorylated level of STAT1 (pSTAT1)/total STAT1 level ratio was done on n = 3 independent experiments. (**b**) Schematic for SMIFH2 treatment. In the SMI-WASH condition (top), fibroblasts were first treated with SMIFH2 (25 μM) at 37°C for 20 min, then washed with PBS to remove unbound SMIFH2, then stimulated with IFN-γ (1000 U/ml) for 10 min. In the SMIFH2 condition (bottom), fibroblasts are always in presence of SMIFH2 (25 μM), then stimulated with IFN-γ (1000 U/ml) for 10 min. (**c**) Western blotting analysis for <u>tyrosine phosphorylation</u> status of STAT1 in cell lysates of fibroblasts in conditions described in (**b**). Quantification was done on n = 5 independent experiments. (**d**) Nuclear translocation status for STAT1 tyrosine phosphorylation reported from immunofluorescence images of fibroblasts with or without SMI-WASH condition (25 μM). White and yellow dashed lines outline whole-cell and nucleus boundaries respectively. The nucleus/cytosol fluorescence ratio for pSTAT1 is quantified from the fluorescence imaging (n = 3 independent experiments). (**e, f**) Immunoblots for tyrosine phosphorylation status of JAK1 (pJAK1) (**e**) and JAK2 (pJAK2) (**f**) in cell lysates of fibroblasts stimulated or not for 10 min with IFN-γ (1000 U/ml) with or without SMI-WASH condition (25 μM) (n = 4 independent experiments). Data shown are mean ± SEM. Statistical significances were determined with an unpaired t-test (**d**) and a one-way ANOVA test (**a,c,e,** and **f**) (∗p < 0.05; ∗∗p < 0.01; ∗∗∗p < 0.001; ∗∗∗∗p < 0.0001; ns, not significant). Scale bar = 20 µm.

### Full length DIAPH3/Dia2 is critical for IFN-γ-induced JAK/STAT signaling

We next sought to identify which specific formin proteins could be involved in IFN-γ induced JAK/STAT signaling. Using siRNA-based screening, we could down-express at a significantly low protein level six individual formin members that all have been reported to be associated with the actin cortex (Breitsprecher and Goode, 2013): FMNL1, FMNL2, DIAPH1 (Dia1), DIAPH3 (Dia2), DAAM1, and FHOD1 (Extended data Fig. 2a). Among those, only Dia2 down-expression was able to significantly inhibit STAT1 phosphorylation (Fig. 2a, Extended data Fig. 2b), and nuclear translocation (Fig. 2b, Extended data Fig. 2c), whereas Dia1, a formin of the same Dia/DIAPH subfamily, had no effect (Fig. 2a, 2b). Furthermore, down-expression of Dia2 decreased to the same extent the activation i.e., the tyrosine phosphorylation levels of JAK1 (Fig. 2c) and JAK2 kinases (Fig. 2d) downstream of IFN-γ stimulation, without modifying the expression of receptor subunits IFN-γR1 and IFN-γR2 at the cell surface (Extended data Fig. 2d, e). Since the JAK/STAT signaling pathway can be triggered by many cytokines and secreted factors, we tested whether Dia2 down-modulation would inhibit JAK/STAT activation in a global manner. Therefore, we compared pSTAT1 level in siDia2 treated fibroblasts upon stimulation by either IFN-γ, IFN-α, or Epidermal Growth Factor (EGF). Remarkably, we observed pSTAT1 decrease only in the case of IFN-γ stimulation. Stimulation by either IFN-α or EGF (Fig. 2e), for which JAK/STAT activation requires endosomal sorting of the activated receptors (Vieira et al., 1996; Marchetti et al., 2006; Zanin et al., 2023) was unaffected by Dia2 down-expression. Altogether, these results demonstrate that Dia2 regulates specifically JAK/STAT signaling at the plasma membrane level in response to IFN-γ.

**Fig. 2.**
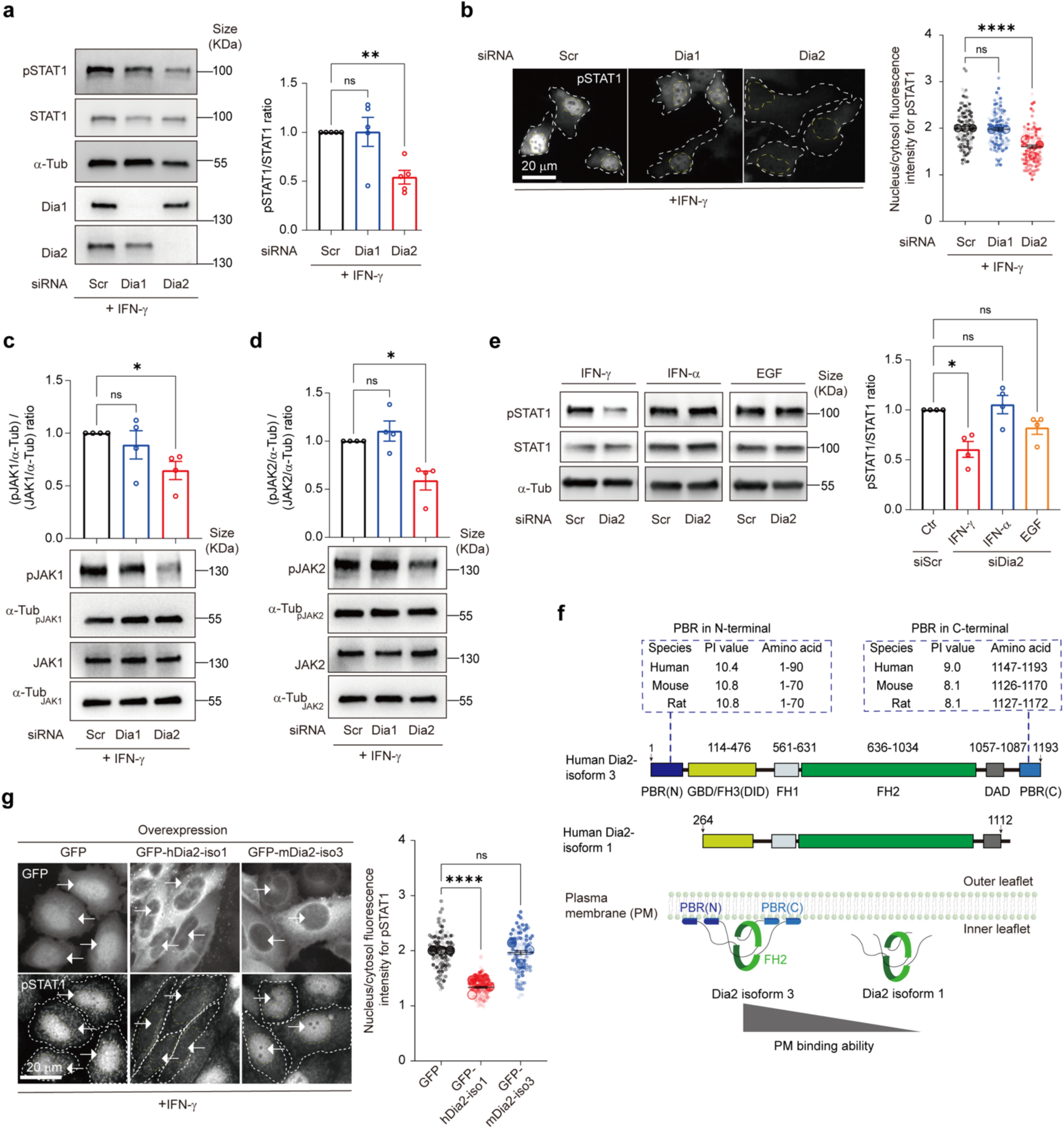
DIAPH3/Dia2 is critical for IFN-γ induced JAK/STAT signaling. (**a**) Immunoblots for pSTAT1 of fibroblasts transfected with non-targeting siRNA (siRNA scramble, siScr), Dia1-targeting siRNA (siDia1) or Dia2-targeting siRNA (siDia2) stimulated for 10 min with IFN-γ (1000 U/ml) (n = 5 independent experiments). (**b**) Immunofluorescence images of pSTAT1 nuclear translocation in fibroblasts transfected with siScr, siDia1 or siDia2 stimulated for 10 min with IFN-γ (1000 U/ml) as described in (**a**). White and yellow dashed lines outline whole-cell and nucleus boundaries respectively. The pSTAT1 nucleus/cytosol fluorescence ratio is quantified from the fluorescence imaging (n = 3 independent experiments). (**c, d**) Immunoblots for pJAK1 (**c**) and pJAK2 (**d**) from cell lysates of fibroblasts transfected with siScr, siDia1 or siDia2 stimulated for 10 min with IFN-γ (1000 U/ml) as described in (**a**) (n = 4 independent experiments). (**e**) Immunoblots for pSTAT1 of fibroblasts stimulated with either IFN-γ (1000 U/ml), IFN-α (1000 U/ml) or EGF (20 ng/ml), transfected with siScr, siDia1 or siDia2 (n = 4 independent experiments). (**f**) Schematic representation of DIAPH3/Dia2 isoforms; PBR (N/C), poly-basic amino acid region in N-terminal/C-terminal; GBD, GTPase-binding domain; FH1, formin homology domain1; FH2, formin homology domain2; DID, diaphanous inhibitory domain; DAD, diaphanous autoregulatory domain. (**g**) Immunoblots for Dia2 and GFP (top). Immunofluorescence images of pSTAT1 nuclear translocation and quantification from fibroblasts transfected with GFP alone (GFP), GFP-mouse Dia2 isoform1 (GFP-mDia2-iso1) or GFP-human Dia2 isoform3 (GFP-hDia2-iso3) (bottom). White and yellow dashed lines outline whole-cell and nucleus boundaries, white arrows indicate cells positive for target overexpression (n = 3 independent experiments). Scale bar = 20 µm. Data shown are mean ± SEM. Statistical significances were determined with a one-way ANOVA test (∗p < 0.05; ∗∗p < 0.01; ∗∗∗∗p < 0.0001; ns, not significant).

In human cells, Dia2 exists in seven isoforms, each with distinct expression patterns. Notably, isoform 1, which lacks both N- and C-terminal parts, is significantly less expressed compared to isoform 7 and the full-length isoform 3 (Stastna et al., 2012). The N- and C-terminal parts of Dia2, which are exclusive to isoform 3, are essential for its proper localization and activation at the plasma membrane (Fig. 2f) (Gorelik et al., 2011; Rousso et al., 2013; Ramalingam et al., 2015; Bucki et al., 2019). The positively charged poly-basic region of Dia2 N-terminal part, referred to as N-PBR, has been shown to bind acidic phospholipids in vitro (Gorelik et al., 2011). Meanwhile, the theoretical isoelectric point (pI) values obtained for both the N-PBR and C-PBR of Dia2 are above 8 (pI > 8), which suggests a potential association with the plasma membrane through electrostatic interactions (Fig. 2f). To understand the role of Dia2 N- and C-domains in the regulation of JAK/STAT signaling, we compared the overexpression of Dia2 full length isoform 3 and isoform1 devoid of N- and C-PBRs. Whereas Dia2-isoform 3 overexpression does not perturb STAT1 activation by IFN-γ stimulation, we observed a drastic inhibition of pSTAT1 nuclear translocation in cells overexpressing the Dia2-isoform 1 (Fig. 2g, Extended data Fig. 2f). Most likely, this inactivation results from the dominant-negative action of the short cytosolic isoform 1, which depletes the substrate from the endogenous full-length isoform 3. As a result, the full-length isoform 3 is unable to process actin filaments at the plasma membrane.

### Dia2 regulation of IFN-γR-dependent JAK/STAT signaling requires RhoA

Rho-GTPases are prominent regulatory factors of actin and myosin dynamics. Various triggers for RhoA activation include the binding of extracellular ligands such as chemokines to G-protein linked receptors (GPCR), cytokines and growth factors to their specific receptors, extracellular matrix (ECM) factors, and intercellular adhesion molecules (ICAM) to integrins (Bros et al., 2019). The formin Dia/DIAPH sub-family (Dia1, Dia2 and Dia3) and Rho-associated protein kinase (ROCK) are the two major downstream effectors of RhoA (Kühn & Geyer, 2014; Lu et al., 2015). The binding of active Rho-GTPases to the N-terminal GTPase binding domain of formins leads to their activation by removing self-inhibition due to internal domains interaction (Breitsprecher and Goode, 2013). But how actin elongation induced by active RhoA is involved in the regulation of plasma membrane receptors is still poorly understood. To determine if Rho-GTPases could regulate IFN-γ-induced JAK/STAT signaling through Dia2 activation, we first analyzed pSTAT1 status upon IFN-γ stimulation in cells treated by Rhosin, a RhoA/C inhibitor (Shang et al., 2012). Rhosin treatment led to a strong decrease in pSTAT1 levels and subsequent nuclear translocation (Fig. 3a-c). Conversely, upon RhoCN03 treatment, a RhoA/B/C activator (Freeman et al., 2018) (Fig. 3a-c), JAK/STAT activation by IFN-γ was fully restored (Fig. 3a-c) confirming the role of Rho-GTPases in IFN-γR function.

**Fig. 3.**
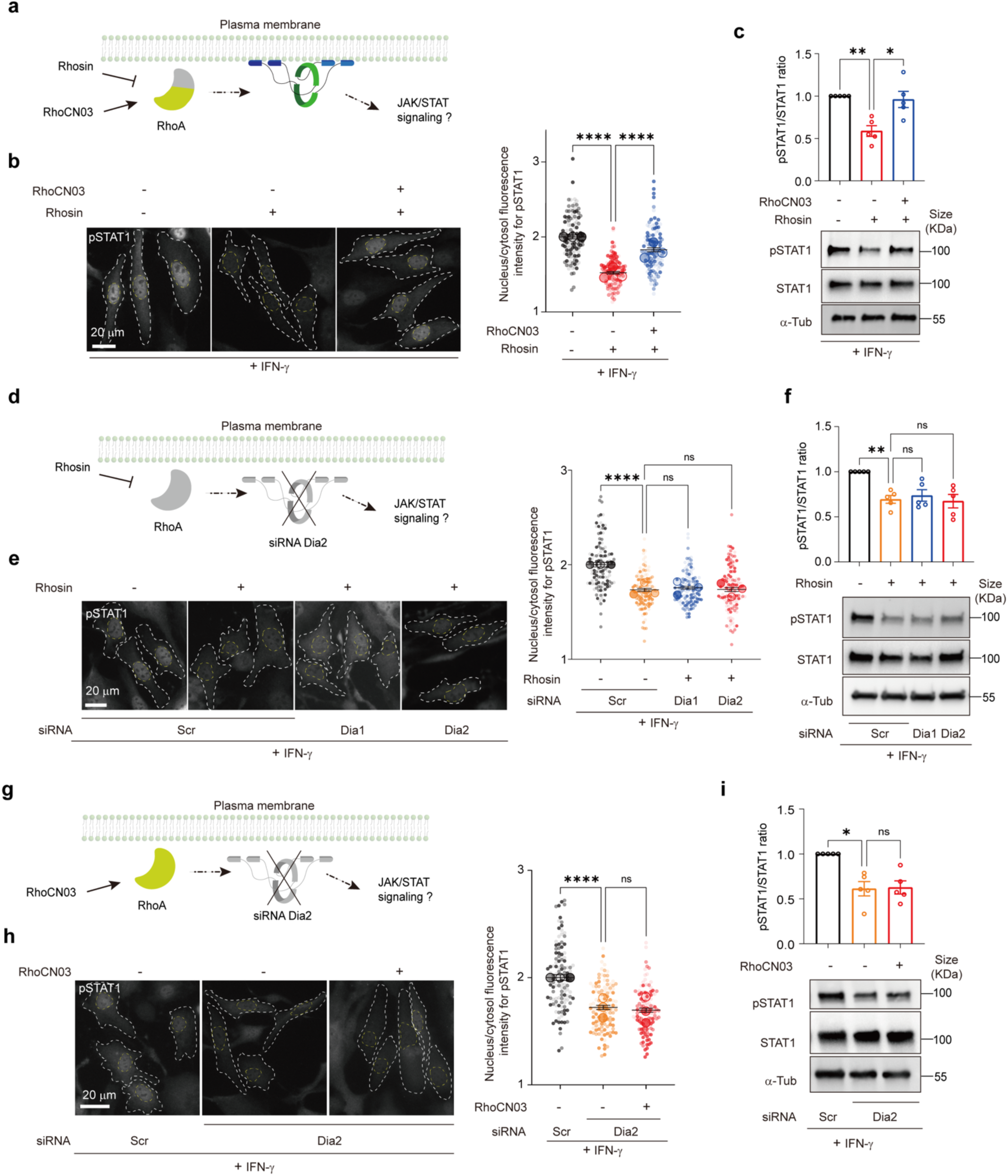
RhoA regulates JAK/STAT signaling through Dia2 activation. (**a**) Schematic representation of the RhoA-regulated Dia2 pathway in presence of Rhosin, RhoA/C inhibitor, and RhoCN03, RhoA/B/C activator. (**b**) Immunoblots for pSTAT1 in fibroblasts in presence of Rhosin (30 μM) for 24 h, then rescue with RhoCN03 (0.25 µg/ml) for 3 h (n = 5 independent experiments). (**c**) Immunofluorescence images and quantification of pSTAT1 nuclear translocation in fibroblasts as described in (**b**) (n = 3 independent experiments). (**d**) Schematic representation of the combination of RhoA inhibition and Dia2 depletion. (**e**) Immunoblots for pSTAT1 of fibroblasts transfected with siScr, siDia1 or siDia2 in presence of Rhosin (30 μM) for 24 h (n = 5 independent experiments). (**f**) Immunofluorescence images and quantification of pSTAT1 nuclear translocation in fibroblasts as described in (e) (n = 3 independent experiments). (**g**) Schematic representation of the combination of RhoA activation and Dia2 depletion. (**h**) Immunoblots for pSTAT1 of fibroblasts transfected with siScr, siDia2 in presence of RhoCN03 (0.25 µg/ml) for 3 h (n = 5 independent experiments). (**i**) Immunofluorescence images and quantification of pSTAT1 nuclear translocation in fibroblasts as described in (**h**) (n = 3 independent experiments). White and yellow dashed lines outline whole-cell and nucleus boundaries respectively. Data shown are mean ± SEM. Statistical significances were determined with a one-way ANOVA test (∗p < 0.05; ∗∗p < 0.01; ∗∗∗∗p < 0.0001; ns, not significant). Scale bar = 20 µm.

To further demonstrate that RhoA regulates JAK/STAT signaling upstream of Dia2, we analyzed STAT1 activation after Rhosin treatment in cells down-expressing either Dia1 or Dia2 (Fig. 3d). Rhosin treatment alone or combined with Dia1 or Dia2 depletion reduced pSTAT1 levels (Fig. 3e) and nuclear translocation (Fig. 3f) to the same extent with no additional inhibition. Since RhoA has many different effectors, it could potentially regulate JAK/STAT signaling through other partners than Dia2. We therefore analyzed STAT1 activation level upon RhoCN03 treatment in Dia2 depleted cells (Fig. 3g) and found that in the absence of Dia2, RhoCN03 can no longer restore pSTAT1 level and nuclear translocation (Fig. 3h,i). Altogether, our data demonstrate that RhoA acts upstream of Dia2 and is necessary for the proper activation of JAK/STAT signaling by IFN-γ.

### Dia2 controls the diffusion dynamics of signaling receptors at the plasma membrane

To investigate the contribution of Dia2 in the regulation of IFN-γR dynamics and signaling, we monitored IFN-γR membrane lateral diffusion in living cells using spot variation fluorescence correlation spectroscopy (svFCS) (Blouin et al., 2016). svFCS is a unique and powerful FCS-based methodology that enables to monitor protein and lipid lateral diffusion regimes at high resolution and with single molecule sensitivity in the plasma membrane of living cells (Lasserre et al., 2008). We carried out svFCS analysis in COS-7 cells stably expressing IFN-γR2-eGFP. At steady state, IFN-γR2-eGFP shows a typical diffusion pattern of dynamic assignment to nanodomains, with relatively high effective diffusion (D_eff_) coefficient and intercept with the time axis (D_eff_ = 0.55 ± 0.08 µm^2^.s^-1^; *t_0_* = 22.93 ± 4.45 ms) (Fig. 4a), as previously described (Blouin et al., 2016). We addressed the potential involvement of formins in the lateral diffusion of IFN-γR2 by first treating cells with SMIFH2. This inhibitor drastically switched IFN-γR2 diffusion from nanodomains to a free-like diffusion behavior with significatively lower D_eff_ and t0 values (D_eff_ = 0.28 ± 0.02 µm^2^.s^-1^; *t_0_* = 10.0 ± 5.14 ms) (Fig. 4a). Similarly to SMIFH2, Dia2 depletion effectively changed diffusion coefficients with a severe impact on the dynamic diffusion of IFN-γR2 at the plasma membrane (D_eff_ = 0.30 ± 0.02 µm^2^.s^-1^; *t_0_* = 8.05 ± 4.18 ms) (Fig. 4a, b). However, Dia1 depletion had only a modest effect on IFN-γR2 lateral diffusion as we could only observe a minor change of *t_0_* and the effective diffusion values (Fig. 4a). This agrees with the lack of effect Dia1 depletion on JAK/STAT signaling activation (Fig. 2a-d). Remarkably, Dia2 depletion led to almost the same *t_0_* value observed with SMIFH2 treatment, confirming that Dia2 is the main formin candidate involved in IFN-γR2 membrane partitioning and function. Moreover, Rhosin treatment modified the dynamic diffusion of IFN-γR2 (D_eff_ = 0.31 ± 0.03 µm^2^.s^-1^), similar to Dia2 depletion (D_eff_ = 0.38 ± 0.07 µm^2^.s^-1^) (Fig. 4b), which strengthens our conclusions regarding the role of RhoA up-stream of Dia2.

**Fig. 4.**
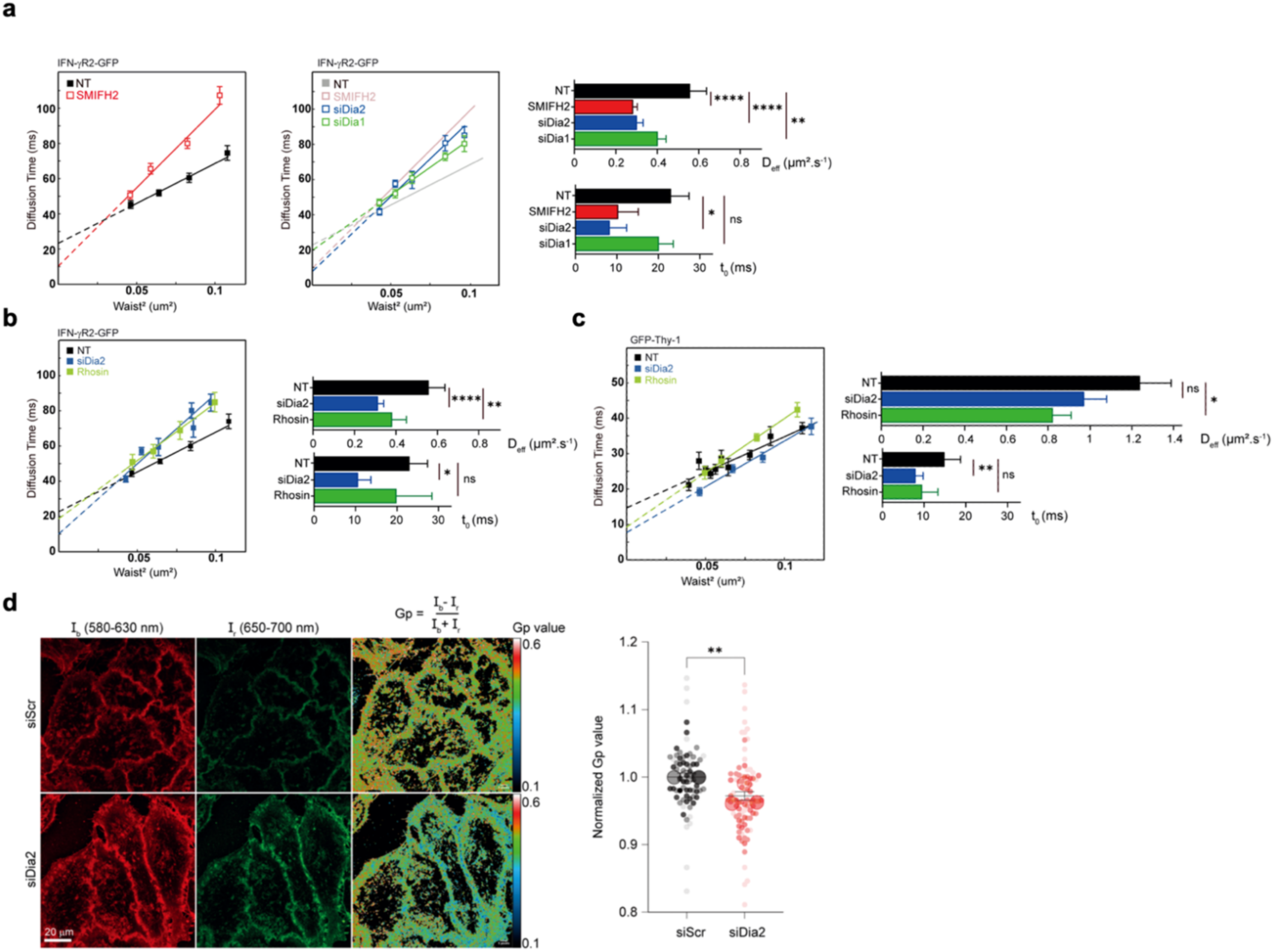
Dia2 controls the diffusion dynamics of membrane receptors at the plasma membrane. (**a**) svFCS measurements of eGFP-tagged IFN-γR2 in COS-7 cells at steady state without (NT) or with SMIFH2 (25 μM) (left), or cells transfected with siDia1 or siDia2 (middle) and histograms of t_0_ and D_eff_ (right). (**b**) svFCS measurements of the eGFP-tagged IFN-γR2 in COS-7 cells at the steady state transfected with siDia2 or treated with Rhosin (30 μM) for 24 h (left part) and histograms of t_0_ and D_eff_ (right). (**c**) svFCS measurements of the eGFP-tagged Thy1 in COS-7 cell at the steady state transfected with siDia2 or treated with Rhosin (30 μM) for 24 h (left) and histograms of t_0_ and D_eff_ (right). For the present svFCS data, t_0_ and D_eff_ are expressed by the mean ± SD, and the statistical significance between two conditions are expressed by a probability of false alarm (PFA) with the threshold defined as follow: ns, PFA >5%; * PFA <5%; ** PFA <1%; *** PFA <0.1%; and **** PFA <0.001%. (**d**) Quantification of GP values at the plasma membrane level of HeLaM cell transfected with siScr or siDia2 (n = 3 independent experiments). Data shown are mean ± SEM. Statistical significances were determined with an unpaired t-test (∗∗p < 0.01). Scale bar = 20 µm.

We also examined the regulation by Dia2 of the lateral diffusion of another plasma membrane receptor associated with lipid nanodomains, the GPI-AP Thy-1 (Lasserre et al., 2008). As observed for IFN-γR2, Dia2 depletion significantly decreases GFP-Thy-1 constrained diffusion (D_eff_ = 1.24 ± 0.15 µm^2^.s^-1^; *t_0_* = 14.78 ± 1.55 ms), transitioning it toward a free-like diffusion pattern (D_eff_ = 0.97 ± 0.11 µm^2^.s^-1^; *t_0_* = 7.82 ± 1.89 ms). This change is again similar to the effects of Rhosin treatment (D_eff_ = 0.82 ± 0,09 µm^2^.s^-1^; *t_0_* = 9.38 ± 2,63 ms) (Fig. 4c). All together, these data suggest that Dia2 regulates the dynamic diffusion of cell surface proteins and receptors.

### Dia2 is required for the formation of cholesterol/sphingomyelin nanodomains at the plasma membrane

Sphingomyelin (SM) is a major sphingolipid present at the plasma membrane of mammalian cells. It is known to form specific membrane lipid nanodomains in conjunction with cholesterol (Chol) and glycosphingolipids (GSLs). Our previous work established the importance of these SM and Chol-dependent nanodomains in the physiological function of IFN-γR (Blouin et al., 2016). Accordingly, a potent SM binding motif was reported in the transmembrane domain of the IFN-γR1 subunit (Contreras et al., 2012). More recently, we have characterized a new Chol binding motif in the IFN-γR2 transmembrane domain, and we found it to be mandatory for IFN-γR activation and signaling (Morana et al., 2022). While the direct relationship between formins and SM-dependent lipid nanodomains has not been established, it has been proposed that short dynamic actin filament structures known as “actin asters”, that are driven by RhoA-dependent acto-myosin contractility, regulate lipid nanodomain formation and receptor activation at the plasma membrane (Kalappurakkal et al., 2019; Wang, et al., 2022). To investigate the potential global impact of Dia2 on the nanoscale organization of lipids at the plasma membrane, we first monitor the behavior of the solvatochromic probe NR12A, which measures lipid ordering (Danylchuk et al., 2019; Carravilla et al., 2021). Using confocal microscopy in live cells, we observed a decrease in the Gp value for the plasma membrane of Dia2 depleted cells compared to control, indicating a more liquid-disordered environment in the absence of Dia2 (Fig. 4d). Therefore, we tested whether Dia2 could participate in the control of lipid nanodomains reorganization at the plasma membrane. Over the years, toxin-derived proteins have been widely used to monitor specific lipids at the plasma membrane under various conditions (Kusumi et al., 2023). Recently, it has become possible to distinguish among the three distinct pools of cholesterol at the plasma membrane: the non-accessible so-called “essential” pool, the accessible pool, and SM-sequestered cholesterol (SM/Chol) pool (Radhakrishnan et al., 2020; Kusumi et al., 2023). We purified and fluorescently labelled Ostreolysin A (OlyA), a toxin known for its specific recognition of SM conformation when associated with cholesterol within SM/Chol complexes (Fig. 5a) (Endapally et al., 2019). In parallel, we also used the Domain 4 (D4 probe) of Perfringolysin O, which only detects the accessible pool of cholesterol (Shimada et al., 2002) (Fig. 5a). In agreement with the literature, OlyA exhibited a concentration-dependent binding on cells, which was abolished upon SM removal by sphingomyelinase (SMase) treatment, confirming its specific affinity for SM (Extended data Fig. 3a,b). It is noteworthy that both SMIFH2 treatment and Dia2 depletion significantly reduced OlyA binding (Fig. 5b,d), indicating a decrease of SM/Chol complexes at the outer leaflet of the plasma membrane. In parallel, both global formin inhibition and Dia2 depletion dramatically increase the surface binding of D4 probe, revealing a higher amount of free cholesterol on the outer leaflet of the plasma membrane (Fig. 5c,e).

**Fig. 5.**
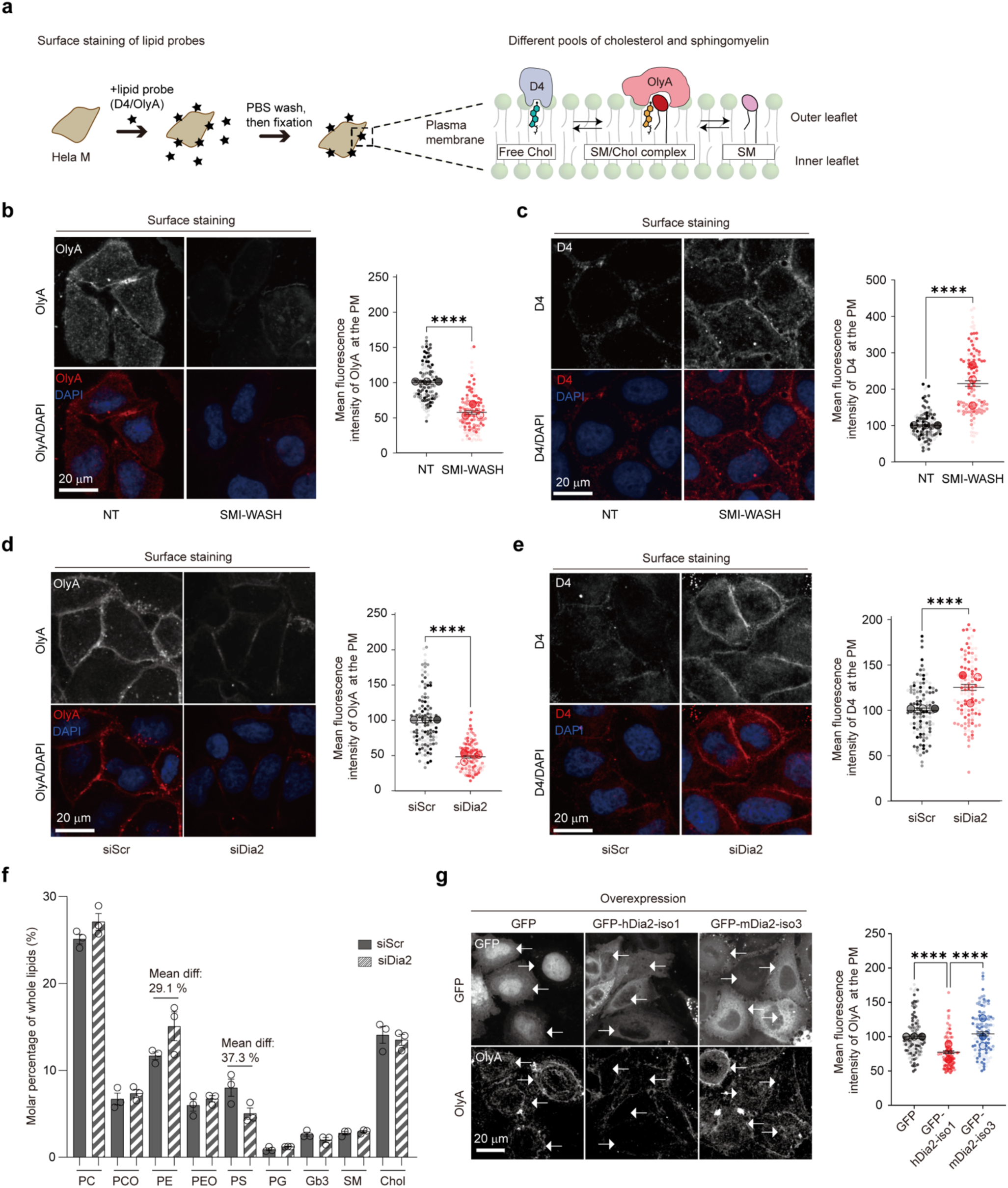
Dia2 regulates SM/Chol complex formation at the plasma membrane. (**a**) Schematic representation of surface staining of the different pools of cholesterol (Chol) and sphingomyelin (SM) in the leaflets of the plasma membrane. HeLaM cells were incubated either with AF647 labelled OlyA (3 μM) or mCherry-D4 (10 µg/ml) at room temperature for 5 min and then quickly fixed. Fungal toxin Ostreolysin A (OlyA) recognizes the SM/Chol complex whereas A domain 4 of bacterial toxin Perfringolysin O (D4) recognizes free Chol, neither complexed with SM nor with membrane receptors. (**b, c**) Surface staining of OlyA (**b**) or D4 (**c**) of HeLaM cell with or without SMI-WASH (25 μM) treatment for 20 min (n = 3 independent experiments). (**d, e**) Surface staining of OlyA (d) or D4 (**e**) in HeLaM cell transfected with siScr or siDia2 (n = 3 independent experiments). (**f**) Lipidomic analysis of total cell lysate from HeLaM cells transfected with siScr or siDia2. The relative abundance of each lipid class was determined by summing the intensities of all molecules in this class and then dividing this value by the total intensities of all identified lipid molecules. Phosphatidylcholine (PC), either phosphatidylcholine (PCO), phosphatidylethanolamine (PE), phosphatidylethanolamine (PEO), phosphatidylserine (PS), phosphatidylglycerol (PG), globotriaosylceramide (Gb3), sphingomyelin (SM), cholesterol (Chol). (**g**) Surface staining of OlyA in HeLaM cells transfected with GFP alone, GFP-hDia2-iso1 or GFP-mDia2-iso3 (n = 3 independent experiments). White arrows indicate positively transfected cells. Data shown are mean ± SEM. Statistical significances were determined with unpaired t-test (**b-e**) and a one-way ANOVA test (**f,g**) (∗∗∗∗p < 0.0001; ns, not significant). Scale bar = 20 µm.

The variations in lipid probes binding observed under Dia2 depletion could not be solely attributed to changes in lipid quantities since lipidomic analysis performed on both control and Dia2-depleted cells, revealed no differences in the total cellular levels of cholesterol and sphingomyelin (Fig. 5f). Moreover, the overexpression of human Dia2 isoform 1 also decreased the SM/Chol complex staining with OlyA (Fig. 5g), in line with its dominant negative effect on IFN-γ-induced JAK/STAT signaling (Fig. 2f,g). In summary, these experiments demonstrate that Dia2 is necessary for maintaining SM/Chol complexes and plays a pivotal role in regulating the equilibrium among the different pool of cholesterol at the plasma membrane.

### Dia2 regulates the nano-organization of SM/Chol complexes at the plasma membrane

We used total internal reflection fluorescence microscopy (TIRFM) coupled to dual-color direct stochastic optical reconstruction microscopy (dual-color dSTORM) coupled to spectral-demixing mode (Lampe et al., 2015; Lehmann et al., 2015) to simultaneously investigate the nanoscale localization of endogenous IFN-γR1 and OlyA (SM/Chol complexes) (Extended data Fig. 4). Dia2-depleted cells exhibited a significant reduction in OlyA labelling density at the plasma membrane compared to control (Fig. 6a and Extended data Fig. 5a) consistent with our observations using spinning-disk microscopy (Fig. 5d). We applied density-based spatial clustering of applications with noise (DBSCAN) cluster analysis (Rahbek-Clemmensen et al., 2017; Nieves et al., 2023) on the STORM coordinates (Fig. 6a). This analysis revealed that in the absence of Dia2, there was a decrease in the fraction of detected OlyA molecules assembled into clusters (Fig. 6b), a reduction in the mean cluster size (Fig. 6c), and a change in the cluster size distribution (Fig. 6e), with individual clusters being more distant from each other (Fig. 6d).

**Fig. 6.**
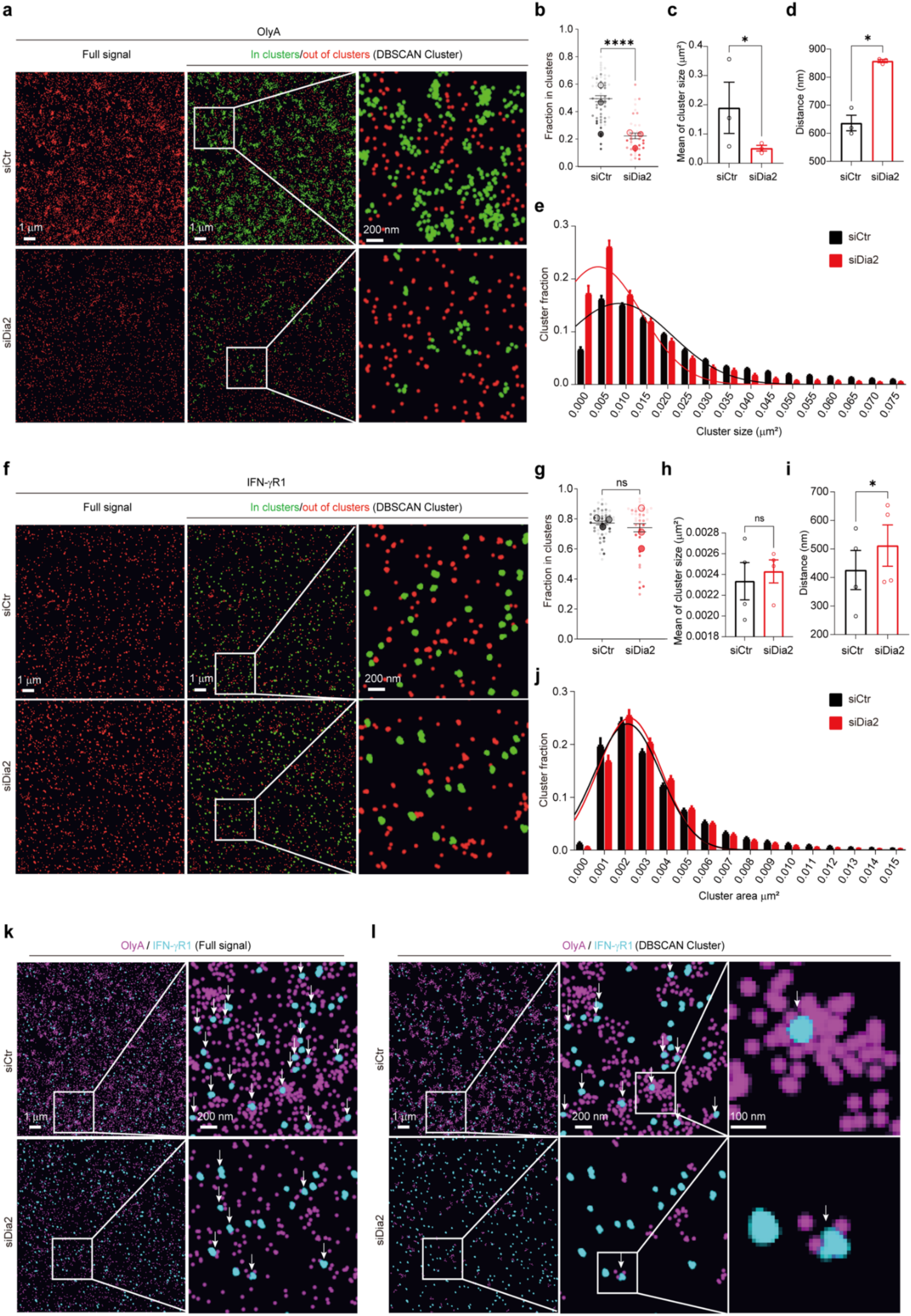
Dia2 controls SM/Chol nanodomain distribution and association with IFN-γR1. (**a-e**) Direct stochastic optical reconstruction microscopy (dSTORM) at total internal reflection fluorescence angles (TIRF) for SM/Chol complex observation at the plasma membrane with AF647 labelled OlyA in HeLaM cells transfected with siScr or siDia2. (**a**) Full signal of OlyA in red color (left panel); DBSCAN-based cluster map (middle and right panels), with molecules within clusters (green) and out of the cluster (red). DBSCAN parameters: ε = 150 nm and MinPts = 5. Representative pictures obtained from white box in Extended data Fig. 4. (**b**). (n = 3 independent experiments). (**b-e**) OlyA data analysis with (**b**) fraction in clusters, (**c**) mean of cluster size, (**d**) distance between clusters and (**e**) cluster size distribution for DBSCAN parameters: ε = 150 nm and MinPts = 5 (n = 3 independent experiments). (**f-j**) TIRF-dSTORM for IFN-γR1 observation at the plasma membrane with IFN-γR1 antibody staining, in HeLaM cell transfected with siScr or siDia2. (**f**) Full signal of IFN-γR1 in red color (left panel); DBSCAN-based cluster map (middle and right panels), with molecules within clusters (green) and out of the cluster (red). DBSCAN parameters: ε = 50 nm and MinPts = 10. Representative pictures obtained from white box in Extended data Fig. 4b. (n = 3 independent experiments). (**g-j**) OlyA data analysis with (**g**) fraction in clusters, (h) mean of cluster size, (**i**) distance between clusters and (**j**) cluster size distribution (n = 3 independent experiments). DBSCAN parameters: ε = 50 and MinPts = 10. (**k, l**) Colocalization analysis of SM/Chol complexes and IFN-γR1 with TIRF-dual color STORM in HeLaM cells transfected with siScr or siDia2. (**k**) Full signal or (**I**) DBSCAN-based cluster map of OlyA (magenta) and IFN-γR1 (cyan), white arrows indicate close association of OlyA and IFN-γR1. DBSCAN parameters: ε = 150 and MinPts = 5 (OlyA), ε = 50 and MinPts = 10 (IFN-γR1). Data shown are mean ± SEM. Statistical significances were determined with a unpaired t-test (**b, g**) and a linear mixed-effects model (**c, d, h** and **i**) (∗p < 0.05; ∗∗∗∗p < 0.0001; ns, not significant).

In contrast, we observed that the propensity of IFN-γR1 to form clusters of the same size was preserved in cells lacking Dia2 compared to control (Fig. 6f-h,j and Extended data Fig. 5b). These data suggest that a decreased amount of SM/Chol complex at the plasma membrane does not alter the formation or maintenance of IFN-γR1 clusters. It is noteworthy that Dia2 depletion resulted in a slight but significant increase in the mean distance between IFN-γR1 clusters (Fig. 6i). We further investigated the potential colocalization between IFN-γR1 and OlyA bound to SM/Chol complex, using their respective coordinates. When considering both the full signal and DBSCAN clustered images at a 20 nm resolution, we observed that IFN-γR1 clusters exhibited reduced proximity with the fewer OlyA clusters in cells lacking Dia2 (Fig. 6k,I, Extended data Fig. 5c). Thus, our data conclusively demonstrate that Dia2 is essential for the nanoscale organization of SM/Chol lipid complexes at the plasma membrane and their association with IFN-γR into lipid nanodomains.

### Dia2 regulates PD-L1 stability at the plasma membrane

We next investigated whether the role of Dia2 in organizing lipids at the plasma membrane could affect other cellular functions beyond JAK/STAT signaling. The programmed cell-death ligand 1 (PD-L1) induces PD-1-mediated T cell exhaustion, inhibiting the antitumor cytotoxic T cell response and leading to tumor survival and growth (Ribas & Wolchok, 2018). Recently, it has been shown that the lipidic environment of the plasma membrane, especially cholesterol, and acidic phospholipids such as phosphatidylserine (PS), is essential for PD-L1 stability and function (Wen et al., 2021; Wang et al., 2022). To assess whether Dia2 influences PD-L1 expression at the plasma membrane, we monitored PD-L1 expression in HeLaM cells by immunofluorescence and western blotting upon EGF and TNF-α stimulations. Both EGF and TNF-α induce PD-L1 expression via the PI3K/AKT and the MEK/ERK pathways, respectively (Jiang et al., 2019). As expected, we observed an induction of PD-L1 expression, with both plasma membrane and total levels increased after 24 h of EGF and TNF-α stimulation (Fig. 7a-c). Under the same conditions, we found that Dia2 depletion decreased both plasma membrane and total PD-L1 levels (Fig. 7a-c). Since we showed that Dia2 depletion does not inhibit EGF signaling (Fig. 2e), these findings strongly suggest that Dia2 regulates PD-L1 stability and degradation through the formation of lipid nanodomain at the plasma membrane. This regulation most likely reflects Dia2 impact on SM/Chol distribution on the cell surface, as well as the cellular levels of PS and PE, as indicated by lipidomic analysis (Fig. 4f).

**Fig. 7.**
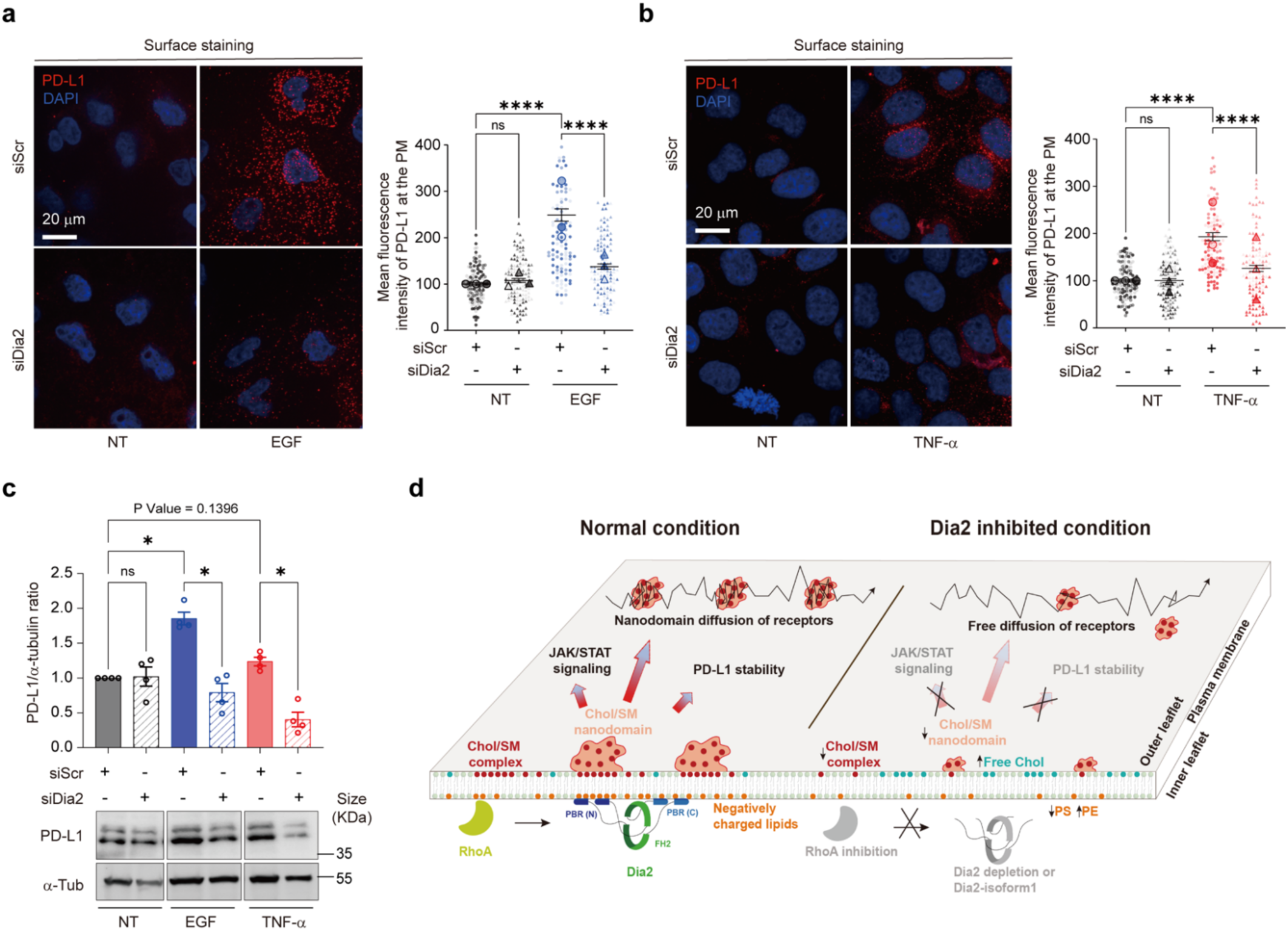
Dia2 is essential for the stability of PD-L1 at the plasma membrane. (**a,b**) Immunofluorescence images and quantifications of PD-L1 surface staining of HeLaM transfected with siScr or siDia2 with or without EGF (**a**) or TNF α (**b**) stimulation for 24 h (n = 3 independent experiments). (**c**) Total cell expression level of PD-L1 analyzed by immunoblots of HeLaM in conditions described in a,b (n = 4 independent experiments). (**d**) Schematic showing how Dia2 controls receptor signaling by organizing plasma membrane lipid (nano)landscape. At steady state, RhoA-Dia2 pathway controls both SM/Chol complex and nanodomain formation at the plasma membrane, which determines plasma membrane nanopartitioning, activation and stability of receptors and lipids. The Dia2-isoform1 that lacks poly-basic amino acid regions loss the SM/Chol complex formation ability, resulting in dysfunction of JAK/STAT signaling. Data shown are mean ± SEM. Statistical significances were determined with a one-way ANOVA test (∗p < 0.05; ∗∗∗∗p < 0.0001; ns, not significant). Scale bar = 20 µm.

## Discussion

The molecular machinery driving or maintaining the assembly of lipid nanodomains around transmembrane receptors is largely unknown. The cortical actin cytoskeleton has been reported to influence membrane organization and molecular diffusion at the plasma membrane (Jaqaman et al., 2011; Fritzsche et al., 2016; Fujiwara et al., 2016; Köster & Mayor, 2016). Based on theoretical and experimental data, it was proposed that actin asters, made of cortical short dynamic actin filaments driven by myosin, regulate the nanopartitioning of membrane proteins (Gowrishankar, et al., 2012). Dynamic actin remodeling has been involved in GPI-AP nanoclustering (Kalappurakkal et al., 2019) and in the formation of CD44-containing actin barriers that build pickets (Freeman et al., 2018; Sil et al., 2020). Actin elongation driven by formins may be involved in these processes, as SMIFH2 treatment affected GPI-APs and CD44 nanoclustering (Kalappurakkal et al., 2019; Sil et al., 2020). How cortical actin and formins regulate the spatio-temporal dynamics of transmembrane protein association with specific lipid nanodomains at the plasma membrane is still elusive (Mayor et al., 2023).

Herein, we investigated the role of formin/Dia2 in lipid nanodomain formation. First, we observed a decrease in global lipid ordering at the plasma membrane in the absence of Dia2, which suggests a partial loss of membrane lipid packing heterogeneity. We then studied the distribution of SM/Chol complexes at the plasma membrane using specific SM/Chol and free cholesterol probes. Dia2 depletion resulted in a significant decrease in the amount of SM/Chol complexes together with a concomitant increase of free cholesterol on the outer leaflet of the plasma membrane. In the absence of Dia2, lipidomics indicate that free cholesterol is released from the SM/Chol complexes since no change was found in the total cellular amount of cholesterol and SM. We studied the distribution of SM/Chol complexes and IFN-γR at the single molecular level with TIRF-dSTORM. We observed fewer and more dispersed lipid clusters at the plasma membrane of Dia2 depleted cells, indicating that the loss of Dia2 prevents the formation of SM/Chol nanodomains. Of note, images from super-resolved microscopy revealed that Dia2 depletion reduced the proximity of IFN-γR1 clusters from SM/Chol complexes. This is consistent with the observations made with svFCS on the loss of dynamic partitioning of IFN-γR in nanodomains. In this context, the presence of a cholesterol binding motif (Sen et al., 2011) and a SM binding motif (Contreras et al., 2012) located at the transmembrane domain (TMD) of IFN-γR1 subunit is informative. More recently, a new consensus Chol-binding motif within the IFN-γR2 TMD, which is required for JAK/STAT signal transduction, has been identified (Morana et al., 2022). The potential interactions mediated by the newly identified lipid binding sites of IFN-γR are certainly critical for inducing conformational changes in IFN-γR upon ligand binding (Blouin et al., 2016). Thus, we propose that Dia2 plays a direct role in controlling the association of IFN-γRs with specific lipids.

An intriguing question is how Dia2, which is active at the inner leaflet of the plasma membrane, can regulate lipid nanodomain formation on the outer leaflet. Outer-leaflet domains enriched in SM and cholesterol, along with specific lipids such as phosphatidylinositol-4,5-bisphosphate (PI(4,5)P_2_) enriched in the inner leaflet, have been reported to mutually regulate each other and to be coupled in a transbilayer manner (Abe et al., 2012, Abe et al., 2021). Such trans-bilayer mechanism could theoretically apply to Dia2 since it can interact with and stabilize PI(4,5)P_2_ domains through the binding of its N-terminal basic domain, a process favored by cholesterol (Bucki et al., 2019). Thus, Dia2 could regulate lipid nanodomain formation not only through actin elongation but also through direct interaction with the plasma membrane. Another mechanism of trans-bilayer coupling was proposed, wherein GPI-APs localized in the outer leaflet interact with the long acyl-chain interdigitations of negatively charged lipid phosphatidyl serine (PS) localized in the inner leaflet. Then, additional membrane associated adapters would couple PS to the dynamic actin-myosin machinery through electrostatic interactions and facilitate the nanoclustering of GPI-APs (Raghupathy et al., 2015). In this context, we observed that Dia2 depletion decreased PS by 37.3%, while increasing by 29.1% the less negatively charged phosphatidylethanolamine (PE) (Fig. 5f). It is therefore possible that the loss of Dia2, which results in the reduction of the overall negative charge at the inner leaflet of the plasma membrane, may perturb the proper recruitment of adapters. The perturbed diffusion of Thy-1 revealed svFCS in the absence of Dia2 (Fig. 4c) may thus reflect the modified clustering of GPI-APs.

We found that RhoA is an essential regulator, upstream of Dia2, for the activation of JAK/STAT signaling by IFN-γ. The spatiotemporal distribution and activation of Rho-GTPases play a significant role in the recruitment and arrangement of actin assembly factors at the plasma membrane. Formins are intrinsically autoinhibited in the cytoplasm and are recruited to the plasma membrane upon interaction with active Rho-GTPases (Pollard, 2016; Chen et al., 2020). Upon RhoA activation, plasma membrane associated-Dia2 can initiate the nucleation and elongation of cortical actin filaments, which in turn could reinforce Dia2 anchorage to the plasma membrane. Remarkably, the overexpression of GFP-Dia2-isoform 1, which lacks N-terminal poly-basic amino acid residues and acts as a dominant negative competitor, led to both the downregulation of SM/Chol complexes at the plasma membrane and the inhibition of JAK/STAT signaling. RhoA itself could directly mediate the formation lipid nanodomains and their assembly around the IFN-γR by interacting with specific lipids and its effectors. This mechanism was revealed for activated Rac1 that partitions into nanoclusters enriched in PI(4,5)P_2_ and PI(3,4,5)P_3_ (Remorino et al., 2017). It is therefore, likely that the active sorting and non-homogenous lateral distribution of RhoA and formins at the plasma membrane play an important role in the formation and maintenance of specific and functional lipid nanodomains.

Finally, when Dia2 is depleted, we observed reduced expression of PD-L1 at the cell surface and intracellularly. A role for formins in PD-L1 expression had never been reported. However, it is known that actin controls the expression of PD-L1 at the plasma membrane, as the actin polymerization inhibitor CytoD significantly reduces PD-L1 protein levels (Miyazawa et al., 2018). Moreover, gene silencing of the ERM scaffolding proteins, namely ezrin, radixin and moesin, that crosslink actin filaments at the plasma membrane, reduces PD-L1 cell surface expression without affecting mRNA levels (Kobori et al., 2021; Tanaka et al., 2021). PD-L1 is a type I transmembrane protein (Zak et al., 2017) presenting a small cytoplasmic domain (CD) encompassing residues 260–290, that is involved in PD-L1 protein stability and degradation. Deletion of the CD tail renders PD-L1 resistant to proteosomal degradation through ubiquitination (Zhang et al., 2018). In addition, palmitoylation of CD Cys272 regulates PD-L1 stability and trafficking (Yang et al., 2019; Yao et al., 2019). PD-L1 degradation is prevented when the CD tail properly binds to the inner leaflet of the plasma membrane through electrostatic interactions (Wen et al., 2021). Two cholesterol-recognition amino acid consensus (CRAC) motifs present in the transmembrane domain of PD-L1 have recently been identified as critical for the protein stability at the plasma membrane (Wang et al., 2022). Based on these studies and the current study, it is highly likely that the strong downregulation of PD-L1 results from increased degradation induced by the absence of a suitable lipidic environment.

The JAK/STAT pathway stands as a paradigm of receptor-mediated signal transduction, with implications in numerous human pathologies. Recently, it has attracted a new surge of interest due to many studies that have revealed the pivotal role of IFN-γ in the intricate process of tumor progression and evasion from immune checkpoints (Benci et al., 2019). Our study has uncovered a significant and novel contribution from the formin Dia2, which plays a crucial role in precisely tuning the JAK/STAT signaling cascade orchestrated by IFN-γR. More broadly, this work brings new mechanistic insights on how actin and lipid nanodomains collaborate to spatiotemporally regulate the nano-organization of membrane receptors and their corresponding bioactivities.

## Acknowledgements

We thank the core facilities and the CurieCoreTech recombinant protein and antibodies platforms of Institut Curie, the staff of the Cell and Tissue Imaging (PICT-IBiSA) and the Nikon Imaging Centre at Institut Curie, as well as the members of the French National Research Infrastructure France-BioImaging (ANR10-INBS-04) for their scientific and technical assistance. We thank Christian Wunder, Julio Lopes Sampaio, Ahmed EI Marjou, Joanna Podkalicka, Christine Viaris De Lesegno, Vincent Fraisier, Feng Ching Tsai, Simli Dey for their help with the lipidomics, OlyA protein purification, fluorescence microscopy and STORM experiments. We thank the following people for providing materials or expertise: Véronique Pizon (Jacques Monod, Paris), Yosuke Senju (Okayama University, Japon), Toshihide Kobayashi (University of Strasbourg, Strasbourg), Andrey Klymchenko (University of Strasbourg, Strasbourg). We thank Patricia Bassereau (Institut Curie, Paris), Ana-Maria Lennon (Institut Curie, Paris), Ludger Johannes (Institut Curie, Paris), Alessandra Cambi (Radboud Institute for Molecular Life Sciences, Netherlands), Xabier Contreras Gomez (Biofisika Institute, Spain) and Guillaume Romet-Lemonne (Institut Jacques Monod, Paris) for comments and suggestions. This work was supported by institutional grants from the Curie Institute, INSERM, CNRS to C.Lamaze, and by specific grants from Agence Nationale de la Recherche (ANR NanoGammaR ANR17-CE15-0032-02 to H.T.H.), Fondation ARC and Ligue Nationale contre le Cancer to C.M.B. C.T.Li was supported by a PhD fellowship from China Scholarship Council (CSC). The Lamaze team are members of Labex CelTisPhyBio ANR-10-LBX-0038, part of the IDEX PSL ANR-10-IDEX-0001-02.

## Author contributions

C.T.Li performed carried out the cell biology, biochemistry, SMI-WASH, siRNA screening and imaging experiments. C.M.B performed pSTAT1 level experiment with different actin relative drug treatment and lipid packing experiments. Y.B., M.D and H.T.H performed and analyzed svFCS experiments. D.M. assisted with dSTORM experiments and analyses. P.G.T. assisted and performed nucleus translocation of pSTAT1 experiment with Dia2-isoforms overexpression. C.T.Li, C.Lamaze and C.M.B wrote the manuscript. C.Lamaze and C.M.B supervised and directed the research. All authors discussed the manuscript and contributed to the preparation of the manuscript.

## Methods

### Cell line, Cell culture

SV40-transformed fibroblasts, HeLaM cells, COS-7 cells were grown at 37 °C under 5% of CO_2_ in DMEM Glutamax (Gibco) with 10% bovine fetal serum. Fibroblasts were supplemented with gentamicin (50 µg/ml) and amphotericin B (1.25 µg/ml).

### DIAPH3/Dia2 constructs

For isoform3 (accession number: Q9NSV4-3) and isoform1 (accession number: Q9NSV4-1), identification and domain numbering are based on entries in the UniProt database. Isoform3 represents the canonical sequence; isoform3 and isoform1 are identical to the major transcript variants deposited in the NCBI nucleotide database, referred to as isoform a (accession number NM_001042517.1/NP_001035982.1) and isoform b (accession number NM_030932.3/NP_112194.2). Isoelectric point (PI) of polybasic regions (PBR) was analyzed with Compute pI/Mw tool of Expasy (Swiss Institute of Bioinformatics, https://web.expasy.org/compute_pi/). All cells were transfected with GFP alone, GFP-human-Dia2-isoform1 (gift from Véronique Pizon) and GFP-mouse-Dia2-isoform3 (gift from Yosuke Senju) using X-tremeGENETM HP DNA Transfection Reagent (Sigma) according to the manufacturer’s instructions cultured for a total of 2 days. Experiments were performed on the validation of overexpression efficiency by immunoblot analysis using GFP and Dia2 antibodies.

### RNA interference-mediated silencing

All cells were transfected with small interfering RNAs (siRNAs) using Lipofectamine™ RNAiMAX Transfection Reagent (Invitrogen) according to the manufacturer’s instructions for 3 days. Experiments were performed on the validation of silencing efficiency by immunoblot analysis using specific antibodies and normalizing to the total level of tubulin used as loading controls. 20 nM of a pool of four siRNA targeting Dia2 were used (J-018997-05, J-018997-06, J-018997-07, and J-018997-07, DharmaconTM), Control siRNA (D-001810-10-05, Dharmacon) and siRNAs targeting other Formins (Dia1, J-010347-06 to −09; FMNL1, J-019176-05 to −08; FMNL2, J-031993-09 to −12; FHOD1, J-013709-09 to −12; DAAM, J-012925-05 to −08; Dharmacon) were used at the same concentration.

### Antibodies

The following primary antibodies were used: anti-pSTAT1 Tyr701 (BD Transduction Laboratories, cat. no. 612132; 1:2,000 for western blotting (WB) and 1:100 for immunofluorescence (IF)); rabbit anti-STAT1 (Cell Signaling Technology, cat. no. 9172; 1:1,000 for WB); rabbit anti-pJAK1 Tyr 1022/1023 (Cell Signaling Technology, cat. no. 3331; 1:1,000 for WB); rabbit anti-JAK1 (Cell Signaling Technology, cat. no. 3332; 1:1,000 for WB); rabbit anti-pJAK2 Tyr 1008 (Cell Signaling Technology, cat. no. 8082; 1:1,000 for WB); rabbit anti-JAK2 (Cell Signaling Technology, cat. no. 3230s; 1:1,000 for WB); mouse anti-IFN-γR1 (BD Biosciences, cat. no. 558935; 1:1,000 for WB and 1:100 for IF); mouse anti-IFN-γR2 (Proteintech, cat. no. 10266-1-AP; 1:100 for IF); rabbit anti-DIAPH1/Dia1 (Cell Signaling Technology, cat. no. 5486s; 1:1,000 for WB); rabbit anti-DIAPH3/Dia2 (Thermo Fisher, cat. no. PA5-109240; 1:1,000 for WB); rabbit anti-DIAPH3/Dia2 (Abcam, cat. no. ab245660; 1:1,000 for WB); rabbit anti-PD-L1 (Extracellular Domain Specific) (D8T4X) (Cell Signaling Technology, cat. no. 86744S; 1:1,00 for IF); mouse anti-human CD274 (B7-H1, PD-L1) (BioLegend, cat. no. 329707; 1:1000 for WB); mouse anti-DAAM1 (Santa Cruz Bio, cat. no. sc-100942; 1:1000 for WB); mouse anti-FHOD1 (Santa Cruz Bio, cat. no. sc-365437; 1:1000 for WB); mouse anti-FMNL1 (Santa Cruz Bio, cat. no. sc-390023; 1:1000 for WB); mouse anti-FMNL2 (Santa Cruz Bio, cat. no. sc-390298; 1:1000 for WB); mouse anti-α-tubulin (clone B512; Sigma, cat. no. T5168; 1:5,000 for WB); rabbit anti-eGFP (Recombinant Antibody Platform, Institut Curie, A-P-R#06; 1:1,000 for WB). The following secondary antibodies were used: mouse-Alexa 488 (Invitrogen, cat. no. A21202); mouse-Cy3 (Jackson ImmunoResearch, cat. no. 715-166-150); mouse-Alexa 647 (Jackson ImmunoResearch, cat. no. 715-606-150); rabbit-Alexa 488 (Invitrogen, cat. no. A21206); rabbit-Cy3 (Jackson ImmunoResearch, cat. no. 111-166-045); rabbit-Alexa 647 (Jackson ImmunoResearch, cat. no. 711-606-152); goat-Cy3 (Jackson ImmunoResearch, cat. no. 705-166-147); goat-Alexa 647 (Jackson ImmunoResearch, cat. no. 705-605-147) were used at 1:100 for IF, 1:5000 for WB; mouse-HRP (Jackson ImmunoResearch, cat. no. 715-035-151) and rabbit-HRP (Jackson ImmunoResearch, cat. no. 711-035-152) were used at 1:5,000 for WB; mouse-CF680 (Sigma, cat. no. Sab4600361) was used at 1:50 for IF.

### Immunoblotting

All cells were lysed in sample buffer (62.5 mM Tris/HCl, pH 6.0, 2% v/v SDS, 10% glycerol v/v, 40 mM dithiothreitol, and 0.03% w/v phenol red). Lysates were analyzed by SDS-PAGE and Western blot analysis and immunoblotted with the indicated primary antibodies and horseradish peroxidase-conjugated secondary antibodies or fluorescently labelled. Chemiluminescence signal was revealed using Pierce™ ECL Western Blotting Substrate, SuperSignal West Dura Extended Duration Substrate or SuperSignal West Femto Substrate (Thermo Scientific Life Technologies). Acquisition and quantification were performed with the Chemi-Doc MP Imaging System (Bio-Rad). The phosphorylated and/or total forms of JAK1, JAK2, PD-L1, IFN-γ cytokine and IFN-γR1 were quantified and normalized to the levels of tubulin in the same lysate. For STAT1, the phosphorylated and total protein levels were assayed on the same blot with the primary antibodies mouse anti-pSTAT1 and rabbit anti-STAT1, and visualized using fluorescence and luminescence, respectively. The ratio of phosphorylated-to-total protein was determined for each time point.

### STAT1 phosphorylation and its nucleus translocation

Cells were cultured in DMEM Glutamax (Gibco) with 10% bovine fetal serum, and then stimulated by switching the medium with DMEM, supplemented with 1000 U/ml of human recombinant IFN-γ (BioLegend), 20 ng/ml of EGF (Sigma) or 1000 U/ml of IFN-α (gift from Biosidus) for 10 min at 37 °C. Prior to IFN-γ stimulation, cells were treated with 25 µM SMIFH2 for 20 min. For the SMI-WASH treatment, cells were treated with 25 µM SMIFH2 for 20 min, following with PBS wash to remove the extracellular SMIFH2 before IFN-γ stimulation. For the Rhosin treatment, cells were cultured in DMEM Glutamax (Gibco) with 10% bovine fetal serum for 24 h, which was then replaced by DMEM Glutamax 0.2% bovine fetal serum, supplemented with 30 µM of Rhosin (Sigma) for 24 h. For the RhoCN03 rescue experiment, Rhosin (30 µM) and RhoCN03 (0.25 µg/ml) (Cytoskeleton, Inc.) were added on cells for 4 h prior to IFN-γ stimulation. For biochemical analysis, cells were washed with ice cold PBS and lysed in SDS Sample Buffer as described earlier. Total lysates were analyzed by SDS-PAGE and Western blot analysis. For the immunoblot signal analysis, pSTAT1 levels were quantified by calculating the ratio between pSTAT1 and STAT1. For immunofluorescent analysis, cells were grown on coverslips, treated as described above and then fixed with cold methanol at −20°C for 10 min. After washing with cold PBS, cells were incubated with primary anti-pSTAT1 antibody for 1 h at room temperature and revealed by using a AlexaFluor488-conjugated goat anti-mouse secondary antibody. pSTAT1 nucleus translocation was quantified with Fiji software (Schindelin et al., 2012) by calculating the nuclei-cytosolic ratio of pSTAT1 signal (Nuclei masks were realized with the DAPI staining).

### Actin staining

To stain for global actin network, cells were grown on coverslips in DMEM Glutamax (Gibco) with 10% bovine fetal serum for 24 h. Cells were then treated with 25 µM SMIFH2 or SMI-WASH at 37 °C for 20 min, and fixed with 4% PFA on ice for 5 min and at room temperature for 20 min. After three PBS washes, cells were permeabilized with saponin (0.2 mg/ml) at room temperature for 10min, then were coincubated with AF647 labeled phalloidin (Cytoskeleton, Inc.) at room temperature for 30 min. Immunofluorescence images were collected by epifluorescence microscopy and quantified with Fiji software (Schindelin et al., 2012) by calculating the mean intensity of fluorophore-labeled phalloidin.

To stain cell cortical actin network. Cells were transfected with siRNAs for 48 h, then transferred to coverslips for 24 h, and fixed with 4% PFA on ice for 5 min and at room temperature for 20 min. After washing with PBS, then directly coincubated with AF647-labeled phalloidin at room temperature for 30 min. Super-resolution images were collected by STORM imaging and analyzed by NEO software (Abbelight).

### Cell surface staining of IFN-γR

Cells were grown on coverslips in DMEM Glutamax (Gibco) with 10% bovine fetal serum, and then washed with ice-cold PBS. Cells were washed with cold PBS, following with primary anti-IFN-γR1 (1:100) (BD Biosciences) and anti-IFN-γR2 (1:50) (Proteintech) antibody on ice for 60 min. After primary antibody binding, cells were washed with cold PBS, then following with 4% PFA on ice for 5 min and at room temperature for 20 min. Cells then were wash with cold PBS again, following co-incubated with fluorophore-labeled secondary antibodies at room temperature for 45 min. Images were acquired using a spinning disk microscope (Inverted Eclipse Ti-E, Nikon; 60x CFI Plan Apo VC oil immersion objective; Spinning disk CSU-X1, Yokogawa; Camera sCMOS Prime 95B, Photometrics). Surface expression levels were quantified using Fiji software (Schindelin et al., 2012) by calculating the mean intensity of fluorescence signal in the sum projected images.

### svFCS

The setup is built around an inverted Zeiss Axiovert 200M microscope, equipped with a C-Apochromat 40× 1.2 NA objective f. A 488 nm excitation laser wavelength illuminates the back aperture of the objective (≈ 2 µW). Once separated from excitation by a 488 nm dichroic mirror and out-of-focus rejection through confocal pinhole, the backward fluorescence is collected by an avalanche photodiode. The back aperture of the objective is under-filled by the laser beam in order to size the FCS volume of observation (Lenne et al., 2006). The defined radius of the excitation beam (namely the waist size, w, ranging from 200-400 nm) is preliminary calibrated according to the Rhodamine 6G diffusion coefficient (280 µm²/s in aqueous solution at 37°C), the estimation of the measurement error being linear with the w² value without exceeding 0.5% even for the larger waist (Billaudeau et al., 2013). For the measurements on living cells, the laser beam focused by the objective is precisely placed at the plasma membrane following a z-scan procedure allowing locating the observation volume at the maximum of fluorescence intensity in the z-axis. The fluorescence signals emitted from the molecules inside the spot volume are then collected. For each observation spot size, each series (between 10 to 15 series of 20 recordings of 5 second each) was performed on different low fluorescent cells (density∼200 molecules/µm² as evaluated from the autocorrelation function (ACF)). For each series, the 20 recordings are selected on the basis of the fundamental FCS assumption in order to keep only the measurements that are in a stationary regime, meaning that the signal fluctuations show a time-invariant profile. Any “abnormal” recordings are discarded; they could be due to strong membrane or cellular movements, or vesicular trafficking. Indeed, the former generates a loose of the focalized spot of observation at the plasma membrane, the later, a collective contribution of individual fluorescently labeled molecules present within endocytic vesicles whose sizes are always smaller than the observation spot. The abrupt and strongly altered signal fluctuation is immediately identifiable on the auto-correlated function due to the invalidation of the time invariance requested for the analysis. The mean of the kept ACFs is fitted with a two-component model.

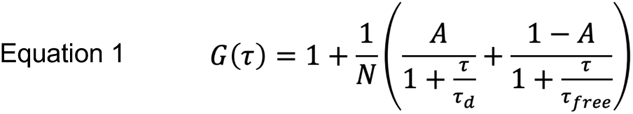

where N is the number of molecules, A the ratio of slow diffusing molecules with a diffusion time τ_d_ (the one used for the diffusion plots) compared to the fast-diffusing ones (presumably corresponding to eGFP proteins non-covalently bound to the receptor). For a given waist, the mean diffusion time calculated from 10 to 15 series is weighted in accordance with the number of kept ACF per series. The estimation of the τ_d_ and w^2^ values were determined using a maximum likelihood estimation considering a Gaussian kernel of the individual and independent recordings with their experimental standard deviations. The diffusion plots were obtained by drawing the average τ_d_ (for each series at a defined waist) and their associated errors (square root of the variance) as a function of w^2^. The values of *t_0_* and D_eff_ were estimated from the diffusion plots by computing the linear regression analysis simultaneously accounting for the estimation errors of τd and w^2^. The estimation variances for *t_0_* and D_eff_ according to equation 2 were given by the Cramer-Rao bound.

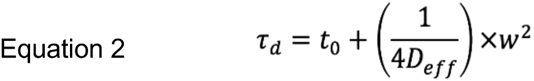

where τ_d_ is the diffusion time, w the spot radius, *t_0_* the y-axis intercept of the diffusion plots, D_eff_ the effective diffusion coefficient proportional to the inverse of the slope of the diffusion plots. The interpretation of the data is done based on the numerical simulations where the variation of *t_0_* corresponds to different modes of membrane compartmentalization (Wawrezinieck et al., 2005; Lenne et al., 2006). In the case of free diffusion (under the Brownian diffusion constraint), the FCS diffusion law projection intercepts the time axis at the origin (*t_0_* = 0); in negative axis (*t_0_* < 0) when molecules are confined by meshwork barriers, or in positive axis (*t_0_* > 0) when molecules dynamically associate with traps and domains (lipid nanodomains) (He & Marguet, 2011; Mailfert et al., 2020).

### Gp value measurement

Lipid packing was monitored with the global polarization (Gp) of NR12A, a solvatochromic photostable plasma membrane-targeting dye (gift from Andrey Klymchenko). Briefly, siRNA transfected HeLaM cells were incubated for 10 min with 200 nM NR12A at room temperature and then wash with DMEM medium without serum. Pictures were acquired with a Nikon A1RHD25 confocal microscope equipped with a CFI Plan Lambda S 100X NA 1.35 silicon immersion objective. Only the images obtained close to the glass substrate, corresponding to the plasma membrane were considered. Gp values for each pixel were then calculated using the plugin for Fiji described in (Carravilla et al., 2021). The following formula was used, where Ib and Ir are the intensities recorded at 580–630 nm (liquid ordered) and 650–700 nm (liquid disordered) channels, respectively:

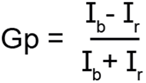

### Sphingomyelin probes purification and labeling

OlyA-His6-WT overexpression was induced at 18°C with 1 mM IPTG for 16 h, and the recombinant protein was purified by nickel chromatography followed by gel filtration (Plateform of protein production, Institut Curie) using the same procedures as described in (Endapally et al., 2019). After purification, OlyA-His6 was labeled with Alexa Fluor 647TM C2-maleimide dye as described previously (Endapally et al., 2019).

### Staining of cell surface lipids with OlyA and D4

For the surface staining, HeLaM cells were transfected with siRNAs for 3 days, SMIFH2 treatment for 20 min, transfected with constructs of Dia2-isoforms for 48 h, or in presence of sphingomyelinase (SMase, Sigma) (0.3 unit/ml) for 1 h. After PBS wash, cells were incubated with either OlyA (3 µM) or D4-mCherry (10 µg/ml, gift from Toshihide Kobayashi) probes in 1% BSA-PBS (v/v) at room temperature for 5 min, then fixed with 4% paraformaldehyde (PFA) for 20 min. Images were acquired using a spinning disk microscope (Inverted Eclipse Ti-E, Nikon; 60x CFI Plan Apo VC oil immersion objective; Spinning disk CSU-X1, Yokogawa; Camera sCMOS Prime 95B, Photometrics). Surface expression levels were quantified using Fiji software (Schindelin et al., 2012) by calculating the mean intensity of fluorescence signal in the sum projected images.

### Sample preparation for lipidomic analysis

HeLaM cells were transfected with siRNA for 3 days, then washed 3 times with 150 mM ammonium bicarbonate (NH_4_HCO_3_) and co-incubated with NH4HCO3 for 5 minutes on ice. After 3 washes by centrifugation with 1 ml of NH_4_HCO_3_ medium, the cell pellets (1.10^6^ cells) were collected with 200 µl of NH_4_HCO_3_ medium, then quickly frozen with liquid nitrogen and stored at −80°C for later use. The lipids were extracted in chloroform/methanol, pre-mixed with internal standard mix, then analyzed at the metabolomics and lipidomics platform of Institut Curie on Q Exactive Plus Hybrid Quadrupole-Orbitrap Mass Spectrometer (Thermo Scientific).

### Sample preparation for dSTORM

Precision Cover Glasses, #1.5H (170 µm thickness) (THOR labs) were cleaned in solution A (125 ml MillQ H_2_O, 25 ml NH_4_OH and 25 ml H_2_O_2_) at 80°C for more than 2 h, then followed by MeOH at room temperature for 10 min. The HeLaM cells (siRNA treatment) are seeded on above cleaned coverslips and cultured for 24 h. The cells were washed with PBS, then directly fixed with 4% PFA on ice for 5 min, then at room temperature for 20 min. The cells followed by an incubation with the SM/Chol probe OlyA (3 µM) and primary anti-IFN-γR1 antibody (1:50) (BD Biosciences) together in 1% BSA-PBS at room temperature for 90 min, then washed by PBS and co-incubated with fluorophore CF680-labeled secondary antibody at room temperature for 60 min. After immunolabeling, a post-fixation step was performed using 4% PFA at room temperature for 15 min after PBS wash. The cells were then quenched for auto-fluorescence from PFA in 50 mM NH_4_Cl at room temperature for 15 min, and then following with PBS wash. The coverslips were kept with PBS until being imaged.

### Acquisition setup of single-molecule localization microscopy dSTORM

Coverslips are mounted on slides with cavity (Sigma) filled with Everspark buffer (protocol here: https://www.idylle-labs.com/everspark-by-eternity#protocol), and then sealed with twinsil (Picodent). Single molecule imaging was done using an Abbelight SAFe360 with the spectral demixing mode (dichroic mirror at 700 nm, Chroma) with two sCMOS cameras (Hamamatsu Fusion BT) mounted on an inverted Nikon Ti2-E microscope with a 100X 1.49 NA objective. The laser used is an Oxxius 640 nm (500 mW, around 3,5 kW/cm² at the sample under experimental condition) at the TIRF angle to specifically excite fluorophores at the plasma membrane near the coverslip (The first few hundred nanometres from coverslip). To summarize the spectral demixing mode, the partially overlapping emissions of the AF647 and CF680 are spectrally split with the 700 nm dichroic beamsplitter before reaching two separated cameras. Then, the intensity ratios between both cameras for each localization are calculated to allocate each particle to the good wavelength (Winterflood et al., 2015). Each acquisition was made with 20 ms exposure time and around 30,000 frames. The data were analyzed with NEO software from Abbelight for noise filtering, cluster fraction, cluster size and localization.

### Density-based spatial clustering of applications with noise (DBSCAN) analysis of dSTORM

We used DBSCAN for cluster analysis. This density-based clustering algorithm searches for clusters by looking for number of localizations within a circle defined by its radius, ε, and its center, p. If the area of the circle contains more than a minimum number of points (MinPts), a new cluster with p as a core object is created. Thus, the DBSCAN algorithm requires the definition of the two input parameters ε and MinPts. The algorithm then subsequently tests all the localizations found in the first cluster for having MinPts within a radius, ε, and assigns them to the cluster, and so forth. If a localization does not have MinPts within ε, this point is defined as a border point of the cluster. We evaluated the MinPts carefully by keeping the real signal and remove the background noise or the monomer fraction localization, and chose MinPts of 5 for OlyA, and MinPts of 10 for IFN-γR1. Then, we evaluated the fraction in cluster, cluster size in a range of ε: 200, 150, 100 and 50 nm. For SM/Chol cluster, we analyzed the fraction in cluster, the cluster size distribution, the mean of cluster size with ε = 150 and MinPts = 5, as 10 times smaller size of probe OlyA (15.6 KDa) than normal antibody. For IFN-γR1 cluster, we analyzed the fraction in cluster, the cluster size distribution with ε = 50 and MinPts = 10, to achieve a search radius that would allow more than one IFN-γR1 protein with antibodies bound next to each other. We analyzed the mean of cluster size with ε = 30 and MinPts = 5, as same result as ε = 50 and MinPts = 10 (data not shown here). We analyzed the distance between clusters of OlyA with ε = 150 and MinPts = 5, and IFN-γR1 with ε = 50 and MinPts = 5. We evaluated the directly localizations with or without DBSCAN between SM/Chol complex and IFN-γR1 at the 20 nm resolution.

### PD-L1 immunostaining

Cells were grown on coverslips in DMEM Glutamax (Gibco) with 10% bovine fetal serum, and then stimulated with EGF (Sigma, 20 ng/ml) or TNF-α (CurieCoreTech recombinant protein - Institut Curie, 20 ng/ml) for 24 h at 37 °C. Cells were washed with ice cold PBS and surface stain with primary anti-PD-L1 (1:50) antibody (BioLegend) on ice for 90 min. After primary antibody binding, cells were washed with ice cold PBS, and fixed with 4% PFA on ice for 5 min and at room temperature for 20 min. After PBS wash, cells were incubated with fluorophore-labeled secondary antibodies at room temperature for 45 min. The images were collected on spinning disk microscope, and surface expression level was quantified with ImageJ software (NIH) by calculating the mean intensity of signal.

### Statistics and reproducibility

All analyses were performed using GraphPad Prism version 8.0 to 9.0, (GraphPad Software). Two-tailed (paired or unpaired) Student’s t-tests and Kolmogorov–Smirnov tests were used when comparing only two conditions. For more than two conditions, the Kruskal–Wallis test was used with Dunn’s multiple comparison test and Tukey’s multiple comparisons test was used. For comparison of Cluster parameters obtained from dSTORM pictures, we used linear mixed-effects model with Jamovi software (https://www.jamovi.org). Significantly different comparisons of means are marked on the graphs with asterisks. Error bars denote s.e.m. or s.d. No statistical method was used to pre-determine sample size. The investigators were not blinded to allocation during experiments and outcome assessment. For comparison between experimental groups (control vs treatment), the samples were assigned randomly to each group. All the control and treated experiments were done on the same day side by side using the same reagents batches. Unless stated otherwise in the figure legends, all representative results shown for immunolabelling, and western blots were performed at least three times independently with similar results. Statistical reports for all cells from the three experiments; smaller solid circles represent individual cells, larger circles represent mean of each experiment, and the same level of transparency corresponds to the same experiment.

---

**Extended data Figure 3.**
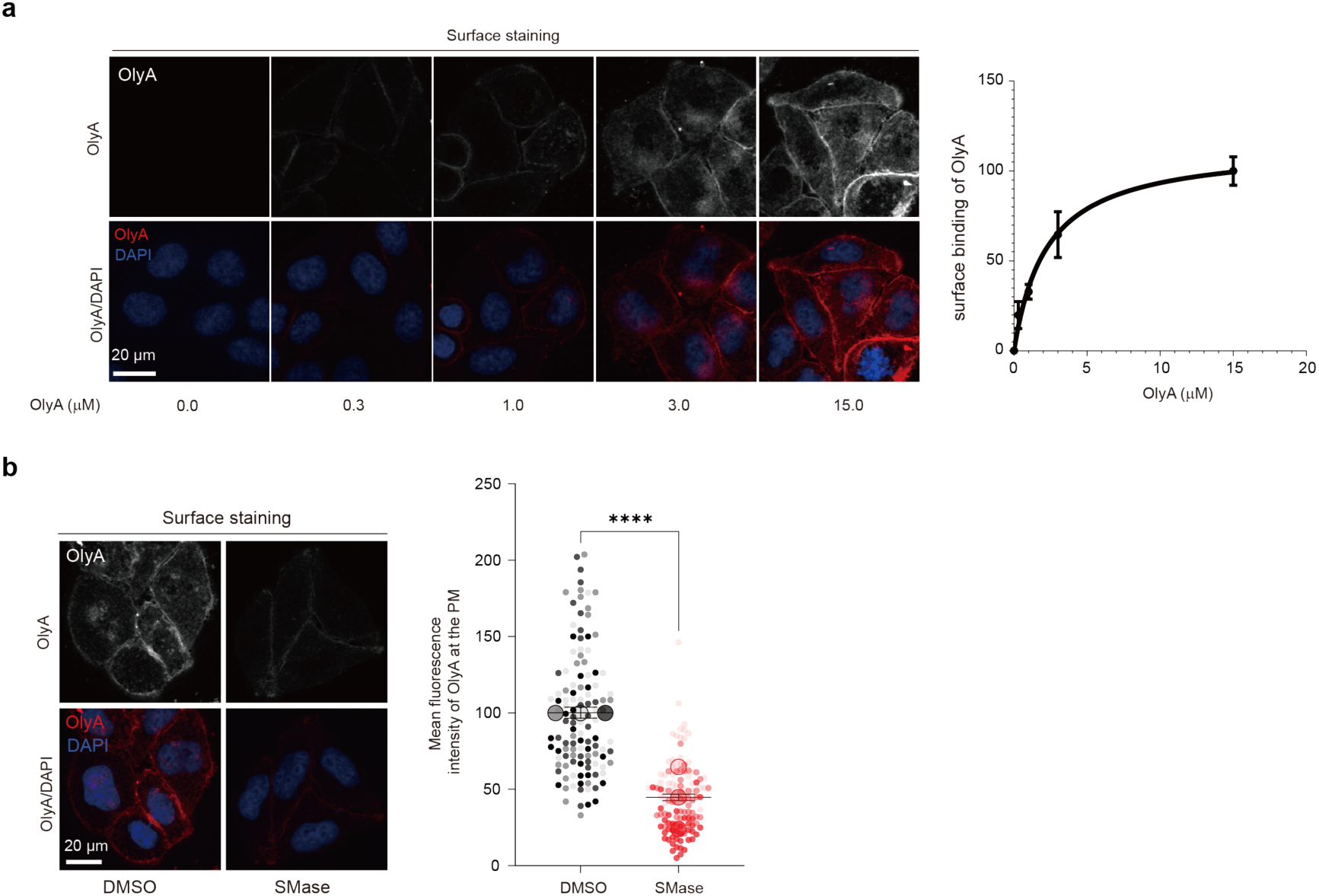
OlyA binding is dependent on sphingomyelin concentration at the plasma membrane. (**a**) Surface staining of OlyA in HeLaM cells at different concentrations. HeLaM cells were incubated with different concentrations of AF647 labelled OlyA at room temperature for 5 min, and quickly fixed (n = 3 independent experiments). (**b**) Confocal images for surface staining of HeLaM cells by OlyA after treatment with Sphingomyelinase (SMase) (0.3 unit/ml) for 1 h (n = 3 independent experiments). Data shown are mean ± SEM. Statistical significances were determined with a paired t-test (∗∗∗∗p < 0.0001; ns, not significant). Scale bar = 20 µm.

**Extended data Figure 4.**
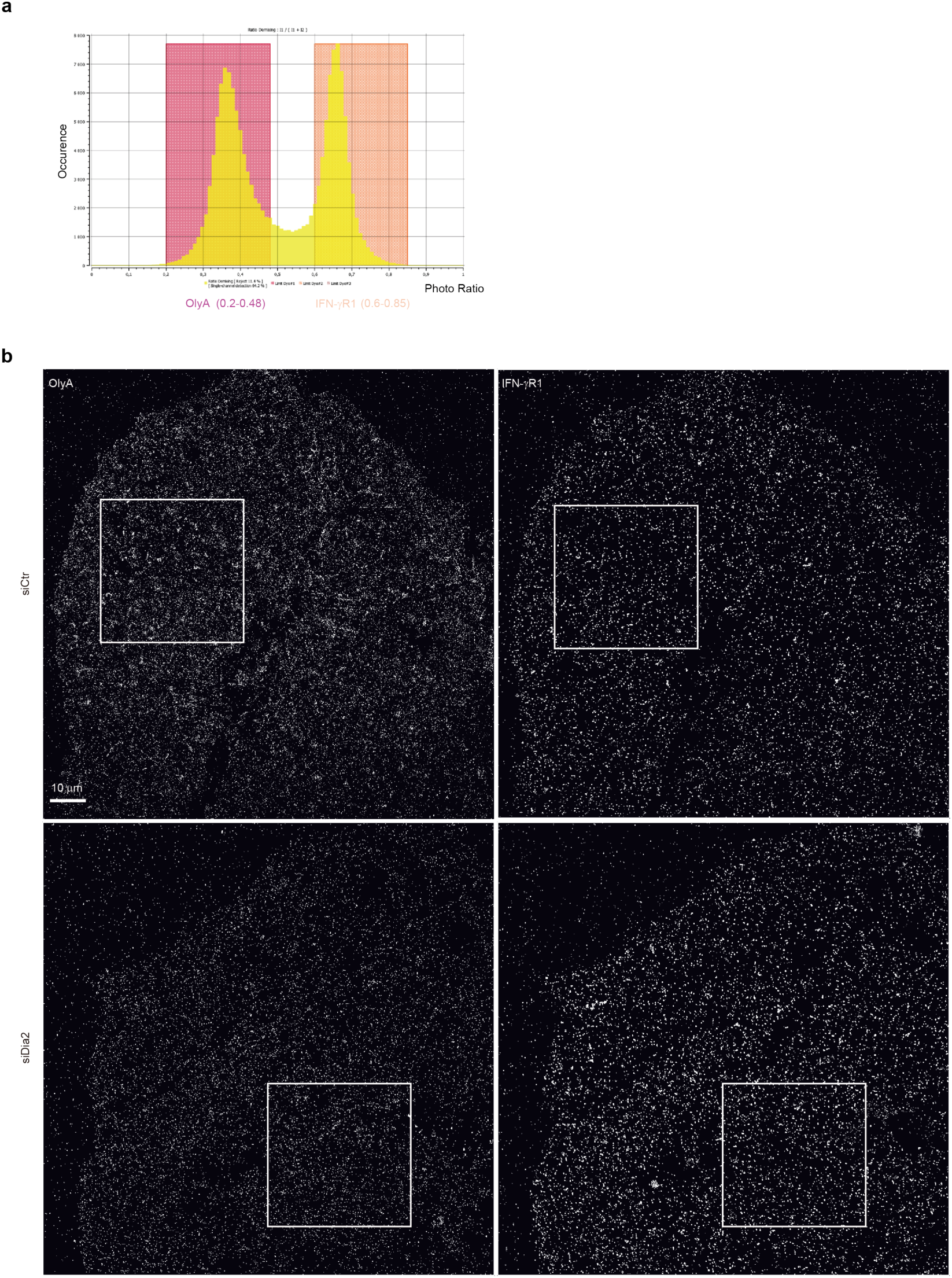
TIRF-dSTORM images of OlyA and IFN-γR1 using a spectral demixing module. (**a**) Demixing photo ratio distribution of OlyA (AF647 signal) and IFN-γR1 (CF680 signal). OlyA signal is collected from 0.2 to 0.48 of photo ratio, and IFN-γR1 is collected from 0.6 to 0.85 of photon ratio. Photon ratio between two cameras placed after a dichroic mirror separating the fluorescence emission from both AF647 and CF680 fluorophores. (**b**) Representative TIRF-dSTORM images for IFN-γR1 and OlyA in HeLaM cells transfected with siSrc or siDia2. White boxes represent zoomed area shown in Fig. 6 and Extended data Fig. 5. Scale bar: 5 µm.

**Extended data Figure 5.**
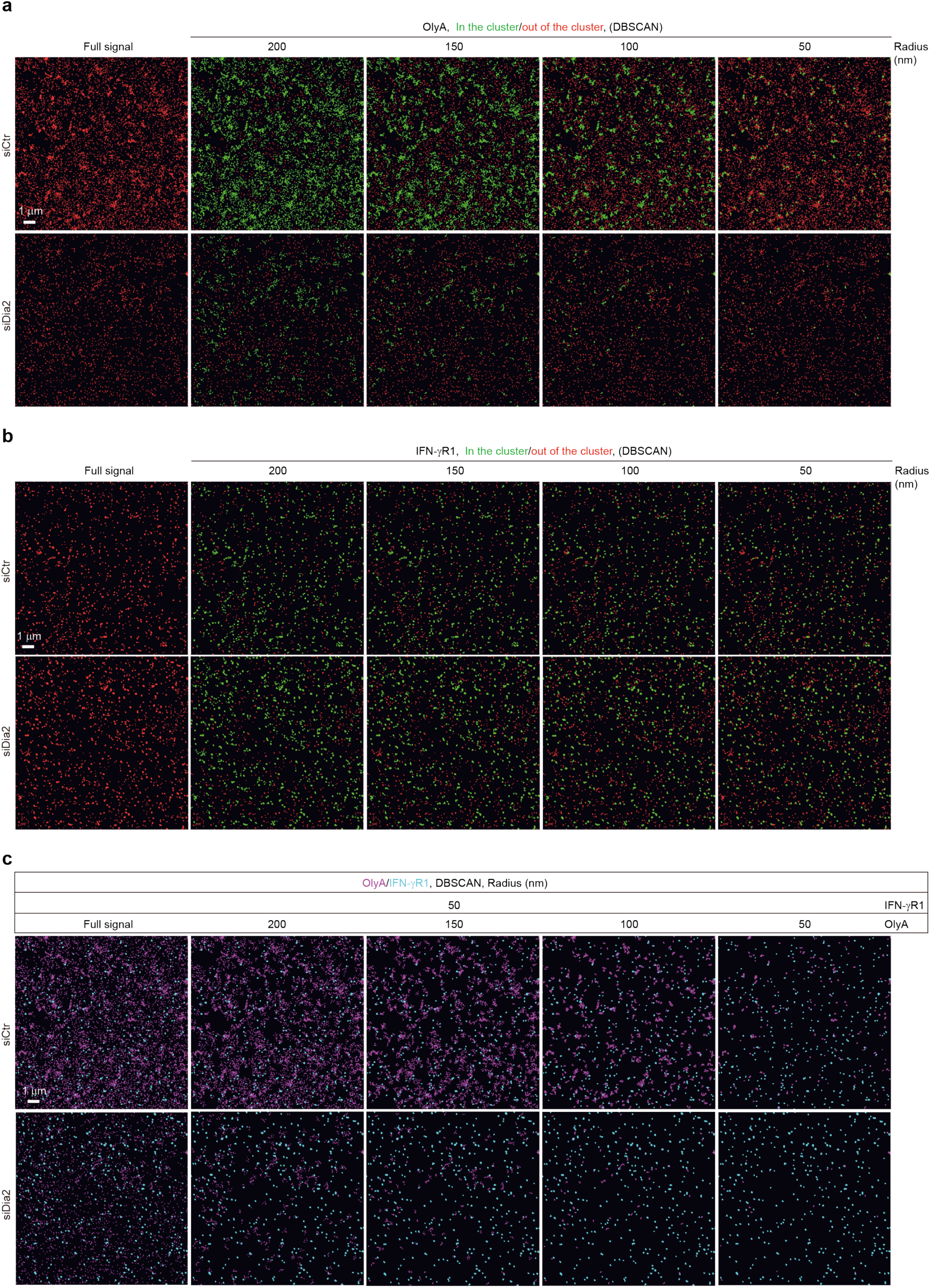
Cluster analysis of SM/Chol and IFN-γR1 using DBSCAN. (**a**) Full signal of OlyA and DBSCAN-based cluster map: within cluster (green) and out of the clusters (red), DBSCAN parameters: MinPts = 5; ε = 200, 150, 100 and 50 nm. Representative images obtained from selected white box in Extended data Fig. 4b. (**b**) Full signal of IFN-γR1 and DBSCAN-based cluster map: within cluster (green) and out of the clusters (red), DBSCAN parameters: MinPts = 10; ε = 200, 150, 100 and 50 nm. Representative images obtained from selected white box in Extended data Fig. 4b. (**c**) Colocalization of SM/Chol complexes and IFN-γR1. Full signal of OlyA (magenta) and IFN-γR1 (cyan), DBSCAN parameters: MinPts = 10 and ε = 50 for IFN-γR1. MinPts = 5; ε = 200, 150, 100 and 50 nm for OlyA. Representative images obtained from merged channel of OlyA and IFN-γR1 shown in (**a**, **b**). Scale bar = 1 µm.

**Extended data Figure 6.**
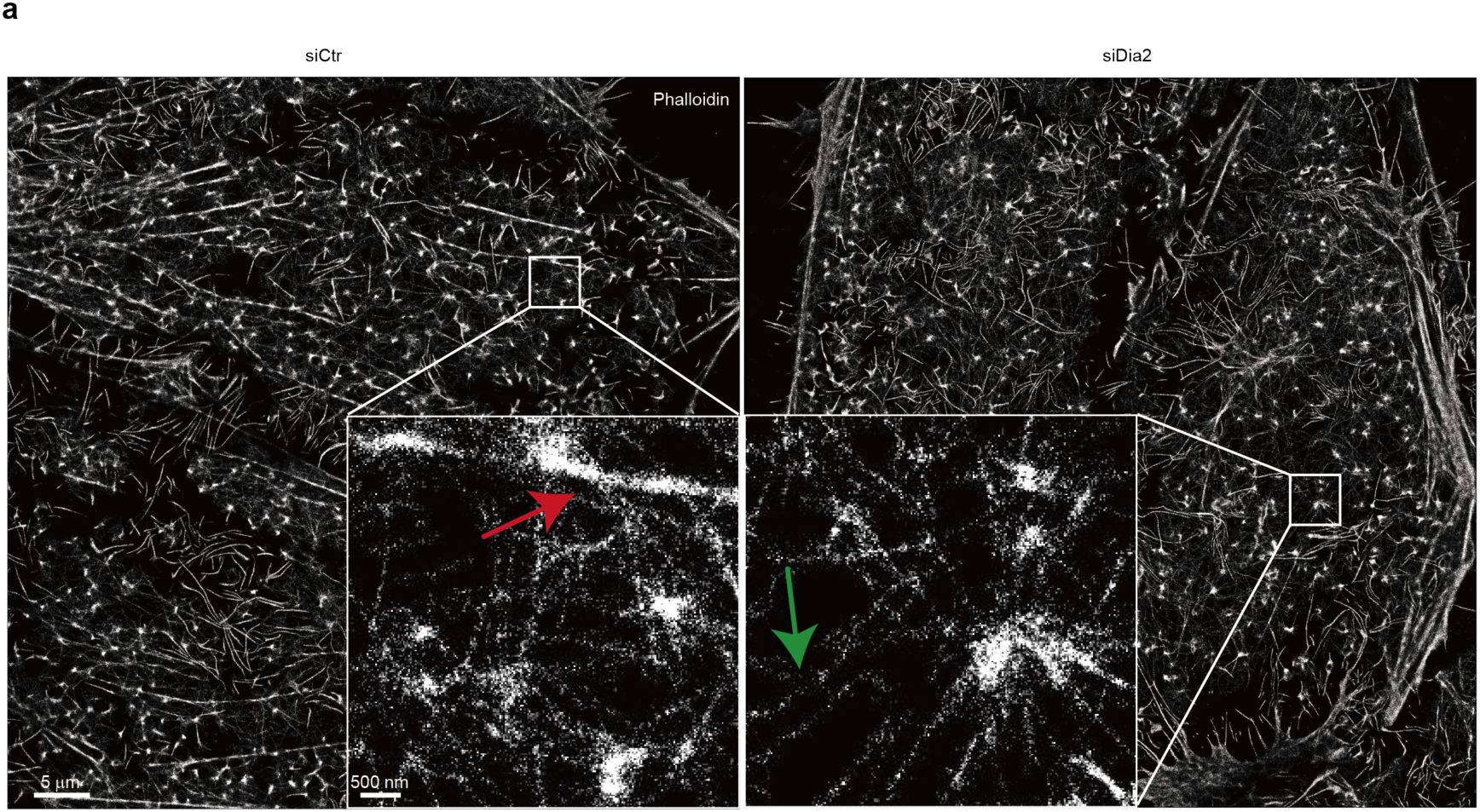
TIRF-dSTORM imaging of the cortical actin network. Scale bar: 5 µm. Magnification of area from select box, scale bar: 500 nm. Red arrow indicates bundled actin filament, green arrow shows the thin unbundled actin filament.

